# Remembering the gist of an event over a lifetime depends on the hippocampus

**DOI:** 10.1101/2021.04.14.439803

**Authors:** Erika Atucha, Shih-Pi Ku, Michael T. Lippert, Magdalena M. Sauvage

## Abstract

A well–accepted view in memory research is that retrieving the gist of a memory over time depends on the cortex, typically the prefrontal cortex, while retrieving its precision relies on the hippocampus. More recent advances indicate that the hippocampal subfield CA1, as opposed to CA3, remains engaged even for retrieving very remote memories and that this engagement coincides with a maximal recruitment of parahippocampal cortical areas (LEC, MEC, PER and POR)^1^. Using a time-window comparable to that used in human long-term memory studies, here we show that CA1 is necessary for retrieving the gist of a memory independently of its age while memory precision specifically depends on CA3 in a time-dependent manner. The precision for the memory of a context-footshock association was tested in mice after one day or very remotely (i.e. after 6 months or one year) allowing for the natural fading of the memory trace. Retrieving recent memories engaged both CA1 and CA3 in control mice as revealed by high levels of RNA of the immediate-early gene *Arc*, strongly tied to synaptic plasticity and memory function. Optogenetic inhibition of CA3 cell firing led to the loss of memory precision, i.e. the retrieval of the gist memory selectively supported by CA1. In contrast, CA1 inhibition abolished memory retrieval and reduced both CA1 and CA3’s activity. At very remote tests, controls retrieved only the gist of the event by recruiting CA1 and parahippocampal areas. Retrieving this gist was selectively abrogated upon CA1 optogenetic inactivation that dramatically reduced parahippocampal activity. Our findings indicate that the hippocampus, specifically CA1, is required for gist memory retrieval even for very remote memories that were previously reported to be hippocampal-independent, while CA3 is necessary for recalling precise memories in a time-dependent manner.

---

The temporally graded retrograde amnesia that affected H.M. after bilateral surgical removal of the hippocampus led to the well-accepted theory of memory consolidation that retrieving recent memories depends on the hippocampus while retrieving remote, less precise, gist memories relies on cortical areas such as the prefrontal cortex^2–6^. This view has been heavily challenged over the past decades as recent advances suggest that the hippocampus is also necessary for retrieving remote memories^7–10^. Specifically, recent reports using methods with higher spatial resolution and studying older memories (> 30 years old in humans^11^ and up to 1 year-old in mice^1^, comparable to 40 years-old memories in humans based on life expectancy) brought compelling evidence of a selective and persistent engagement of CA1, even for the retrieval of the most remote memories. In addition, parahippocampal areas (the PER, POR, MEC and LEC) were also found to be maximally recruited for the retrieval of the oldest memories, much like what is expected from the prefrontal cortex^1,12–16^. Since memories become less precise over time^5,17^, the ongoing engagement of CA1 is indicative of an important role of CA1 in the retrieval of gist-like memories despite the well-accepted concept of hippocampal independency of the retrieval of remote memories. Conversely, the hippocampal subfield CA3 is no longer recruited for retrieving very remote memories^1^. This together with the fact that computational theories predict a crucial role of CA3 for pattern completion (the completion of full memory representations based on detailed features of these representations^18)^ suggest a pivotal and time-limited role of CA3 in retrieving memory precision. The extent to which CA1, CA3 and the parahippocampal areas contribute to retrieving the gist of memories and their precision is however not well-understood.

Dissociating the contribution of CA1, CA3 and that of parahippocampal areas to memory retrieval over time is beyond the standard (3T) spatial resolution of fMRI studies in humans. In contrast, animal models are especially well-suited for this type of studies because they allow for imaging brain activity with cellular resolution and transient and targeted subregional brain manipulations. Here, we investigate the role of CA1, CA3 and parahippocampal areas in the retrieval of precise and gist memories and examine the neural activity underlying these processes at the system level. The precision of the memory for a context-footshock association was assessed behaviorally in mice following conditioning using a time-window comparable to that used in humans for studying remote memory: one day, 6 months and 1 year after memory formation (comparable to investigating 20 and 40 years-old memories in humans based on life expectancy^1^). To this end, mice were exposed to the context in which they previously received a footshock and subsequently, to a similar but safe context^19,20^ (Fig. 1A). CA1 or CA3 function were altered by inhibiting cell firing upon memory retrieval using the inhibitory Archeorhodospin (ArchT)^15,21^ (green stars in Fig. 1A,B). The neural substrates engaged during the retrieval of the context-footshock association were assessed in ArchT expressing mice and their controls using an imaging technique yielding cellular spatial resolution based on the detection of the RNA of the immediate-early gene *Arc*, closely tight to synaptic plasticity and memory function^22–27^ (Fig. 1B).

**Fig.1.**
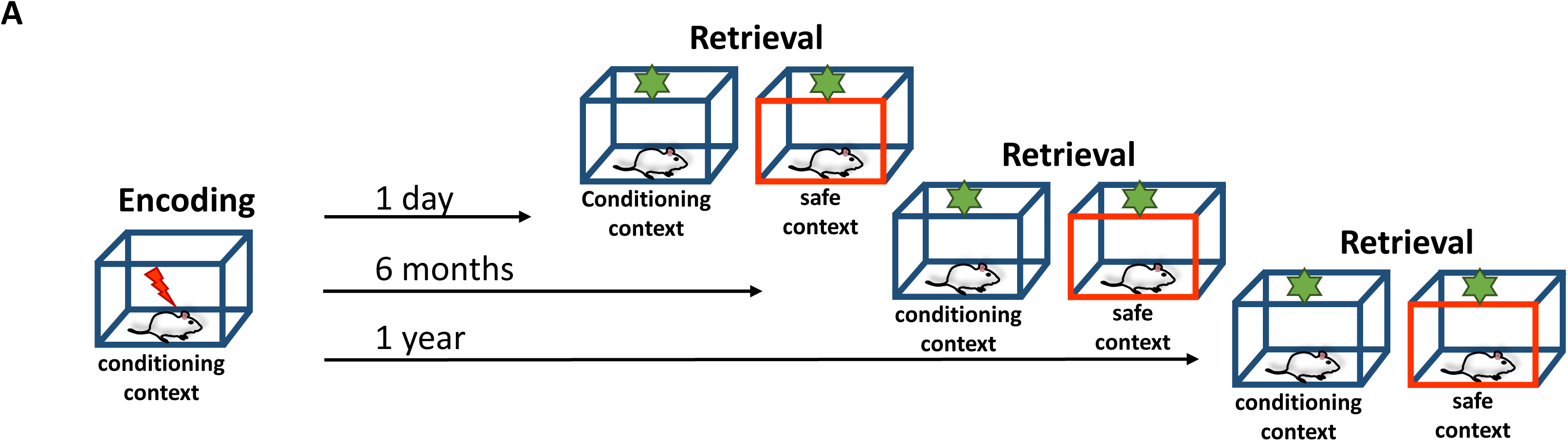
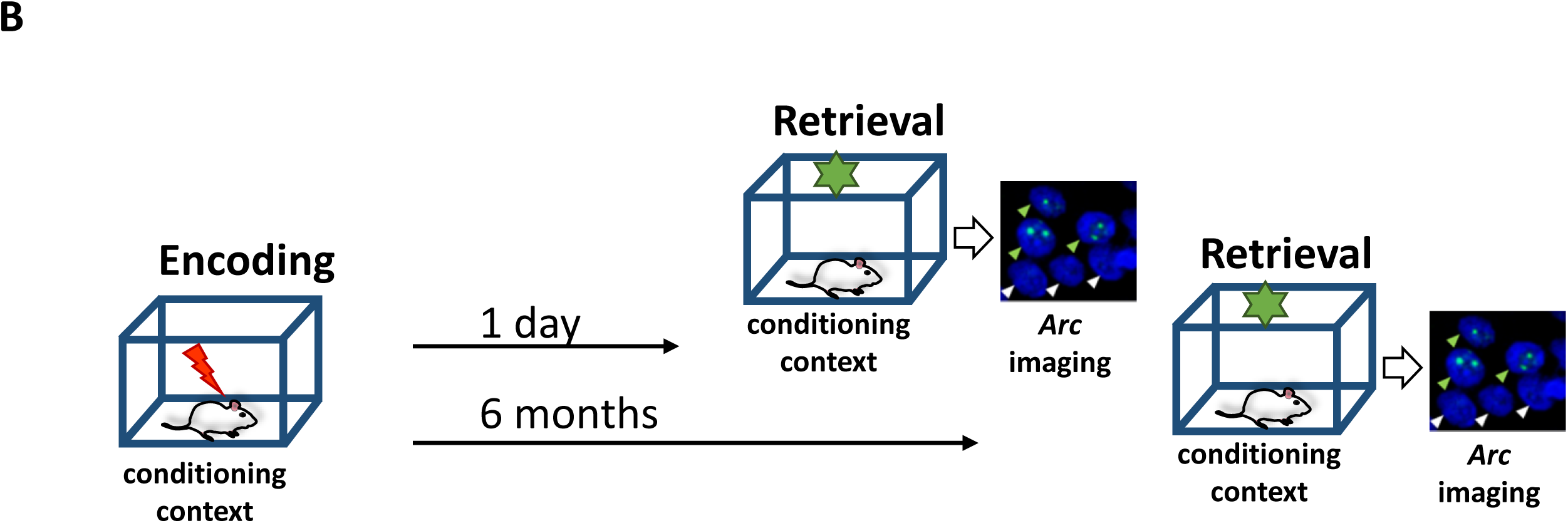
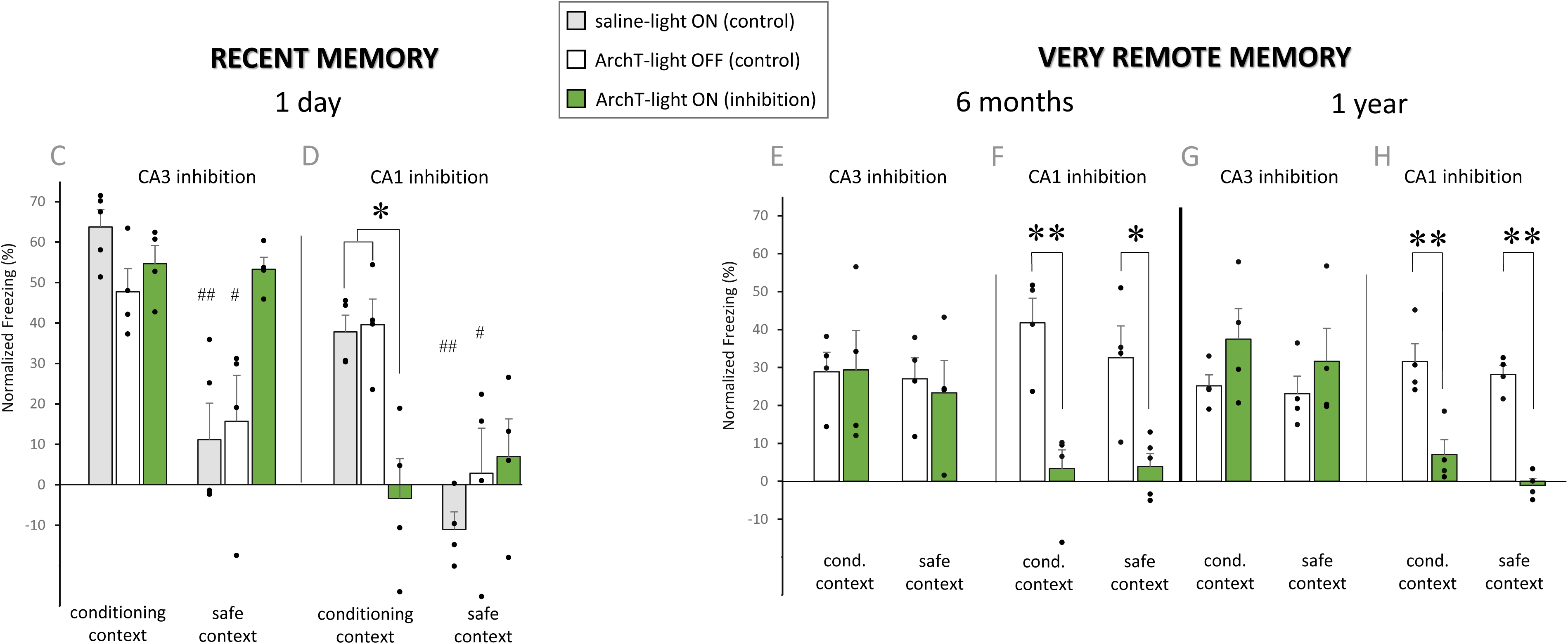
Effects of selective and transient cell firing inhibition in CA1 or CA3 on the retrieval of precise and gist contextual fear conditioning memory over half a lifetime. *A. Assessment of memory performance.* During encoding, mice explored freely the conditioning context for 2.5 min upon which a single 1 mA footshock was delivered (red thunder). Mice remained 2.5 min in this context before being returned to their homecage. During the retrieval phase, the precision of the memory for the context-footshock association was tested for recent (1 day-old) and very remote memories (6 months and 1 year-old) by re-exposing mice to the conditioning context and consecutively to a similar but safe context (in which no shock had been delivered) for 5 min each. Selective freezing to the conditioning context is indicative of the retrieval of a precise memory whereas freezing to both contexts is indicative of contextual generalization, i.e. indicative of the retrieval of an imprecise gist-like memory. Bilateral cell firing inhibition in CA1 or CA3 was implemented by stimulating the inhibitory opsin ArchT with a LED for the first half of each retrieval test (green stars). *B. Imaging memory retrieval over half a lifetime.* Patterns of activity induced during the retrieval of the context-footshock association were imaged upon re-exposure to the conditioning context for recent (1 day) and very remote (6 month-old) memories by detecting the pre-mRNA of the immediate-early gene *Arc* (green arrowheads: examples of CA1 *Arc* positive cells; white arrowheads: *Arc* negative cells; cell nuclei are labeled in blue with DAPI)*. C-H) Effect of the transient optogenetic inactivation of CA1 or CA3 on memory retrieval over time.* Mean ± SEM of normalized freezing levels of conditioned mice at the recent (1 day) and very remote (6 months and 1 year) retrieval tests. Dot plots are overlaid (n=4-5 mice per groups). *C)* Memory precision was lost upon CA3 inhibition when the memory was recent (green bars compared with grey or white within context and green bars compared between contexts) whereas remote gist memory retrieval was unaffected by CA3 inhibition when the context-footshock memory had decayed 6 months and 1 year after conditioning in controls (*E* and *G,* same comparisons). In stark contrast, interfering with CA1 function, dramatically impaired the retrieval of the memory when only the gist of the memory was retrieved (i.e. when very remote memories were retrieved; *F* and *H:* green bars compared with white) as well as when memory was recent (*D*: green bars compared with grey, white). * P<0.05, **P<0.01 for comparisons between inhibition and control groups: # P<0.05, ## P<0.01, for comparisons between conditioning and safe context (see Extended Data Fig. 1 A-E for additional control measures for the task performance).

At recent memory test (1day), controls retrieved a precise memory for the context-footshock association independently of the hippocampal subfield targeted (CA1 and CA3; Fig. 1C,D). This was indicated by a selective freezing behavior to the conditioning context, reflecting a successful discrimination between the context in which the footshock was delivered (the conditioning context) and the safe context (# compare grey or white bars between conditioning and safe context: paired t-tests: t_3-4_ > 3.22, P<0.038, one sample t-test against 0: conditioning contexts: *P*< 0.001; safe contexts: *P*> 0.264; Fig 1C,D). This was no longer the case at very remote memory tests in line with reports showing a loss of memory precision over time in studies focusing on early remote memories^5^ (i.e.1 month-old). Indeed, controls froze to a comparable extent upon exposure to conditioning and safe context, indicative of the generalization of the fear memory to the safe context and the retrieval of the gist of the memory six months and one year after encoding (compare white bars between conditioning and safe context: paired t-tests: t_3_<0.665, P>0.554, one sample t-tests against 0: conditioning and safe contexts: *P<* 0.01: Fig 1E-H). Note that no advert effects of the light delivery *per-se* were observed on behavioral performance as freezing levels to the conditioning or the safe context were comparable between saline-light ON and ArchT-light OFF control groups (compare grey with white bars: conditioning context: both t_6-7_<1.362, p<0.215, safe context: both t_6_<1.167, P>0.288; Fig 1 C,D). Also, since this potential ‘light’ effect was not expected to depend on the age of the memory but on delivering the light, the saline/light ON control group was generated only for recent memories according to the 3Rs principle (Replacement, Reduction and Refinement).

Importantly, an additional control experiment showed that, independently of delay imposed between conditioning and retrieval phases, mice that did not receive a footshock during the encoding phase of the task (i.e. the no-shock groups) explored to a comparable extent the conditioning and the safe context (both F_2,9_<2.53; P>0.135 for context _X_ memory age interaction effects; Extended Data Fig.1A &B). Hence, differences in freezing or neural activity levels reported in the present study are unlikely to stem from differences in non-mnemonic parameters such as locomotion, motivation to explore contexts or context preference. Furthermore, a separate control experiment, showed that exposing conditioned mice to the conditioning context prior to exposure to the safe context did not affect the freezing levels in the safe context (unpaired t-test: t_7_=0,332, P= 0.43; Extended Data Fig. 1E).

Imaging neural activity for the retrieval of the precise context-footshock association in controls at the recent memory test revealed that among all medial temporal lobe areas (MTL) studied only CA1 and CA3 hippocampal subfields were recruited (Fig 2A,C; white bars: CA1 or CA3 vs other MTL areas: all P<0.001 Tukey post-hoc tests; F_5,18_>18,221; P<0.001 area effect; one sample t-tests against 0: CA1 and CA3: P<0.013, parahippocampal areas: P> 0.22, but MEC P = 0.059 for CA3 inhibition; see Fig2E for a graphic summary). This result is in line with both the system consolidation and the multiple trace theories predicting the engagement of the hippocampus during the retrieval of recent memories, albeit without dissociating the involvement of CA1 from that of CA3^4,6,7,9^. In addition, it confirms more recent findings according to which parahippocampal areas do not play a crucial role in retrieving memories when they are recent^1^. By contrast, imaging brain activity in controls at the very remote memory tests (when controls recall the gist of the memory and no longer its precision showed that parahippocampal areas were recruited at least as much as CA1 and that CA3 was disengaged (white bars in Fig 2 B,D; see Fig.2H for a graphic summary: parahippocampal areas vs CA1: all P>0.^14^3 Tukey post hoc tests; CA3 vs other areas: all P<0.0010 Dunnet’s post hoc tests; F_5,18_>15.624, P<0.001 area effects; one sample t-tests against 0: CA3: P>0.068 for CA3 inhibition, P= 0.012 (negative values due to normalization) for CA1 inhibition, all other areas P < 0.040). Of note, since each CA1 and CA3 cell firing inhibition had a similar effect on memory retrieval 6 months or 1 year after conditioning (compare white with green bars between Fig 1E and G and between Fig. 1F and H: both F_1,24_<0.0001, P>0.99 lack of memory age x inhibition x context interaction effects), neural activity was only imaged 6 months after encoding. Importantly, it was also established that differences in patterns of activity between recent and very remote memory tests were unlikely to stem from differences in the strength of the memory retrieved one day and 6 months after encoding as memory performance of control groups (ArchT/light OFF) from which brain activity was imaged was comparable between recent and very remote memories (compare grey bars: t_6_<1.005, P>0.354, unpaired t-tests; Extended data Fig.3A&B). Note that similar results were found for the ArchT/light OFF control groups used to measure memory precision (compare white bars in conditioning context between Fig. 1C and E and between Fig. 1D and G: t_6_<1.563; P>0.16 unpaired t-tests). Altogether, these results show that, in controls, CA1 and CA3 are recruited during the retrieval of the memory for the precise context-footshock association in controls while CA1 and parahippocampal areas are engaged for the retrieval of very remote gist memories (see Fig 2E and H for a graphic summary).

**Fig.2.**
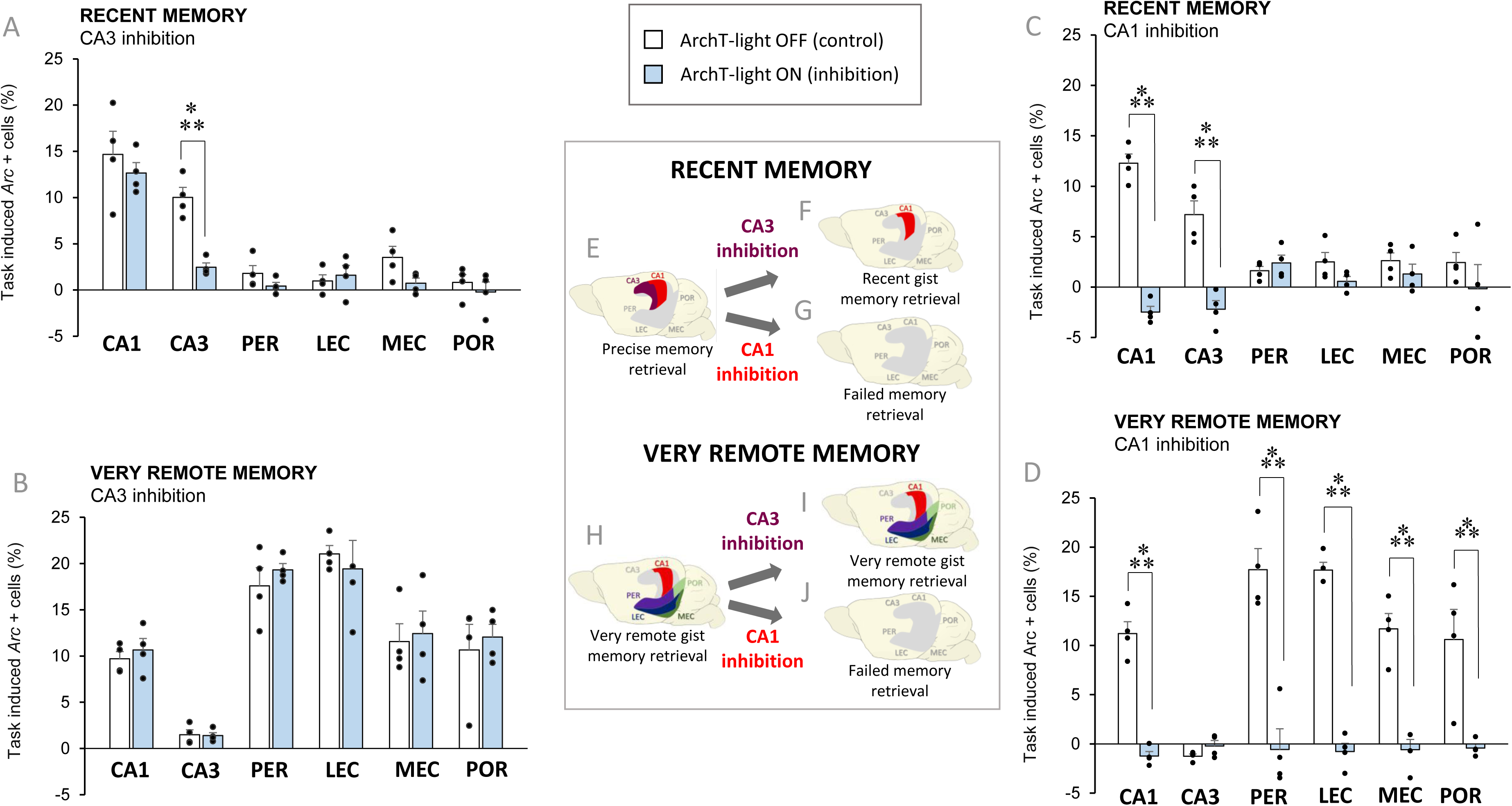
Effects of the transient inactivation of CA1 or CA3 on hippocampal and parahippocampal neural activity during memory retrieval. Mean ± SEM of normalized percent of *Arc* positive cells in conditioned mice (n=4 per group). Dot plots are overlaid. *A*) Inactivation of CA3 that led to the retrieval of the gist of the recent memory, i.e. the loss of memory precision selectively reduced CA3’s engagement during memory retrieval while CA1 remained engaged (compare blue with white bars; see Fig. 2E,F for a graphic summary). This indicates that CA3 is necessary for the retrieval of memory precision, but not for that of memory gist and conversely, that CA1 contributes to retrieving the gist of recent memories. In contrast, compromising CA3 function did not affect neuronal activity in areas recruited during the retrieval of very remote gist memories (i.e. CA1 and parahippocampal areas’, Fig. 2B) confirming that CA3 is not required for gist memory retrieval independently of the age of the memory trace, that its role in memory precision is time-limited and that CA1 participates in retrieving memory gist even when memories are very remote. In addition, inhibiting cell firing in CA1 revealed a causal relationship between CA1’s neural activity and gist memory retrieval as neural activity in CA1 was abolished when very remote gist memory retrieval failed (Fig 2D; see Fig 2H,J for a graphic summary). This effect was however more global than in the case of CA3’s inhibition as in addition to CA1, all areas recruited during very remote memory retrieval, i.e. the parahippocampal areas, were also affected (Fig. 2D). A similar network effect was observed at recent memory test on areas engaged during recent memory retrieval (i.e. CA1 and CA3; Fig2 C). *Graphic summary* of the effects of the transient inactivation of CA1 or CA3 on MTL neural activity and the nature of the recent (E-G) and very remote (H-J) memory representations retrieved. ***P<0.001.

To test the causal relationship between CA1 or CA3 function and the retrieval of precise and gist-like memories, cell firing in CA1 or CA3 was transiently inactivated optogenetically by stimulating the inhibitory opsin ArchT expressed in either CA1 or CA3 (see Extended Data Fig.2 for *in-vivo* electrophysiological evidence of ArchT–induced cell firing inhibition (A, B), ArchT expressing virus spread (C, D) and optic fiber stubs location (E,F)). In addition, the effect of these manipulations on memory performance and MTL neural activity was assessed. At recent memory tests, i.e. when controls retrieved the precise contextual fear memory, inhibition of CA3 cell firing led to the generalization of the fear memory to the safe context. Indeed, freezing levels were high and comparable between the conditioning and safe context (green bars: unpaired t-test: P=0.806; one sample t-test against 0: both P<0.001; Fig 1C), indicating that CA3 inhibition abolished the retrieval of the precision of the memory and promoted the retrieval of the gist of the event. Specifically, CA3 inhibition enhanced the freezing level in the safe context compared to control groups (t_6-7_>3.199; P<0.0186 unpaired t-tests) without affecting freezing in the conditioning context (t_6-7_<1.534; P>0.169 unpaired t-tests; Fig 1C). Hence, the results of optogenetic inhibition of CA3 at recent memory test are twofold: CA3 is necessary for retrieving precise memory as mice undergoing CA3 cell firing inhibition fail to discriminate between conditioning and safe context and CA3 is not critical for retrieving gist-like memories as CA3 inhibition promotes the retrieval of these memories. The selectivity of CA3’s contribution to memory precision was further supported by the imaging results revealing that brain activity was reduced only in CA3 upon CA3 cell firing inhibition at recent memory test (compare white with blue bars: CA3: P< 0.001, other MTL areas: all *P* > 0.684 Tukey post hoc tests; F_5,36_=3.093, P = 0.020 for area x inhibition interaction effect; Fig. 2A). In addition, these data hinted at a selective role of CA1 in retrieving the gist of recent memories as only CA1 remained engaged upon CA3 cell firing inhibition that led to gist memory retrieval (blue bars: CA1 vs all areas: P<0.001, all others P > 0.676 Dunnet’s tests; F _1,36_=12,583; P=0.001 for area effect; one sample t-test against 0: CA1: *P* = 0.001; parahippocampal areas: *P* > 0.210; Fig 2A; see Fig.2E, F for a graphic summary). A direct causal link between CA1 and gist memory retrieval is established at a later point.

The lack of involvement of CA3 in gist memory retrieval was further supported by behavioral findings at the remote memory tests, when only the gist of the memory was retrieved (Fig.1E,G). Indeed, cell firing inhibition of CA3 6 months and one year after conditioning did not significantly affect memory performance in either context (compare white with green bars: F_1,16_<0.412, P>0.532 lack of inhibition x context interaction effects) demonstrating that CA3 does not contribute in a decisive manner to gist memory retrieval even when memory traces are very remote. This behavioral finding is in agreement with the disengagement of CA3 observed for the retrieval of very remote memories in control mice in the present study (white bars, Fig 2B, D) that had also been observed in a previous study of ours that did not focused on investigating the nature of the memory representations retrieved^1^. As expected in such a case, altering CA3 function during the retrieval of very remote memories did not affect CA3’s level of activity nor that of any other areas recruited during very remote gist memory retrieval. As such, CA1 and parahippocampal areas remained highly engaged in the group undergoing inhibition (blue bars: one sample t-tests against 0 for CA1 and other areas P<0.017; since 2-way ANOVA showed F_5,36_=1.883; P=0.122 lack of inhibition x area interaction, CA3 was no longer recruited: P=0.715; Fig.2B, see Fig.2H,I for a graphic summary). Thus, transiently inhibiting cell firing in CA3 reveals a selective role of CA3 in the retrieval of the precision of recent memories and a lack of a contribution to gist-like memory retrieval independently of the age of the memory trace. Moreover, this result suggested a meaningful role of CA1 in the retrieval of gist like memory over time.

To directly test the causal relationship between the recruitment of CA1 and the retrieval of precise or gist-like memories, CA1 cell firing was transiently inhibited. At recent memory test (Fig 1D), CA1 cell firing inhibition abolished the retrieval of the context-footshock association (compare grey or white bars with the green bar: conditioning context: *P*< 0.0102 unpaired t-tests; green bar: one sample t-test against 0: P=0.755) without affecting freezing levels to the safe context (compare grey or white bars with the green bar: t_7_<1.362; *P* > 0.215 unpaired t-tests; green bar: one sample t-test against 0: *P* = 0.327). These results are symptomatic of a critical role of CA1 in the retrieval of recent memories but not in retrieving its precision as, in contrast to CA3’s manipulation, CA1 cell inhibition did not promote context generalization. Failure to retrieve recent memories upon CA1 cell firing inhibition was paralleled by a marked drop in neural activity not only in the targeted area (CA1) but also in CA3, while parahippocampal areas’ activity remained negligible (compare white with blue bars: CA1, CA3: *P*< 0.012; parahippocampal areas: *P*>0.115 Tukey post hoc tests; F_5,36_=15.040, P=0.001 for area x inhibition interaction effect; blue bars: one sample t-test against 0: all P>0.075; F _1,24_=0.966; P=0.335 as area effect; Fig.2C, see Fig.2E,G for a graphic summary). In summary, imaging results indicate a network effect of CA1 inhibition on the recruitment of the hippocampus during recent memory retrieval as opposed to a local effect observed upon CA3 inhibition.

A similar picture unfolded at remote memory tests as inhibition of CA1 cell firing also abrogated the retrieval of memory 6 months and one year after conditioning independently of the context studied (compare white with green bars: all P< 0.011; both F _1,6-7_> 30.240, P=0.001 for inhibition effects; Fig 1F,H). Importantly, at very remote tests CA1 inhibition abolished the retrieval of generalized memories as controls retrieved at this time point only the gist of the memory and no longer precise representations. This effect was mediated by the disengagement of all MTL areas supporting the retrieval of very remote gist-like memory in controls (CA1 and the parahippocampal areas) while it did not affect CA3’s which was not recruited at this time point (compare white with blue bars: CA3: P = 0.141, all other areas: P< 0.012 Tukey post-hoc tests; F_5,36_= 11.936, P=0.001 for area x inhibition interaction effect; blue bars: one sample t-test against 0: P > 0.252; Fig.2D, see Fig.2H,J for a graphic summary). Thus, these results indicate that, also at very remote memory tests, inhibiting CA1 cell firing during memory retrieval affects not only CA1 but the entire MTL network involved in retrieving gist-like memories. This finding is complementary to earlier results of ours according to which CA1 is not engaged in control mice failing to retrieve very remote (one year-old) memories but did not focus on identifying the nature of the memory representation retrieved^1^. In summary, these results extend and confirm the findings that had emerged from the selective inhibition of CA3’s cell firing reported in the present manuscript according to which CA1 plays a crucial role in the retrieval of the gist of memories independently of the age of the memory trace and does not play a decisive role in retrieving their precision.

Extensive evidence shows that the hippocampus is crucial for retrieving precise memories^3–6,8,28–30^ and that cortical areas are increasingly involved over time in supporting the retrieval of the gist of these memories, as retrieving the former requires intact hippocampal function while retrieving the latter is dramatically altered by insults to the neocortex^12,15,16^. We show now that the hippocampus, specifically the hippocampal subfield CA1, is necessary for the retrieval of the gist of memories and that the contribution of the hippocampus to memory precision and memory gist retrieval is subfield-dependent. How does the hippocampus contribute to the retrieval of precise memories and their gist as the memory ages at the system level? We recently showed that CA3 was recruited to a similar extent for the retrieval of recent and early remote memories (1day and 1 month-old, respectively)^1^. In addition, computational studies have suggested a key role of CA3 in memory precision retrieval via pattern completion^18^. Given the present results, we propose that CA3 might contribute to the retrieval of memory precision by relying on pattern completion and that this role is restricted to the most recent memories (i.e. up to one month old memories) as CA3’s disengagement for the retrieval of very remote memories coincides with the retrieval of memories that are no longer precise, i.e. gist-like memories. This together with the fact that detailed features of memories degrade as memory traces age, rendering them less precise^5^, make it plausible that pattern completion might no longer be a resource for the retrieval of the most remote memories. Indeed, detailed features might then be so degraded that they can no longer be used as cues for completing full memory representations. In contrast CA1, which remains engaged even for the retrieval of very remote memories would play a major role in retrieving the gist of these memories together with parahippocampal areas, maximally recruited at this time point, as interfering with CA1 function at this time point abolishes the retrieval of gist memory and is paralleled by a sharp decrease in parahippocampal areas’ recruitment. This hypothesis is supported by earlier findings reporting a clear disengagement of CA1 and reduced parahippocampal activity in mice failing to retrieve such remote memories^1^in a study that did not test for causality. This, together with the fact that CA1 remains the only MTL area engaged for retrieving recent gist memories (i.e. when CA3 function is compromised), suggest that CA1 plays a pivotal and selective role in retrieving the gist of memories independently of their age and plays a preponderant role in retrieving the most remote gist memories with the support of parahippocampal areas

Overall, these findings are crucial because they demonstrate that the retrieval of the gist of remote memories requires specifically the engagement of the hippocampal subfield CA1 while gist memory retrieval has been to date considered as relying on cortical reactivation^4,5,8,9^. Furthermore, our results bring the first evidence that parahippocampal areas might play a role comparable to that of the PFC in memory consolidation, at least for the retrieval of the most remote memories. Further investigations will be necessary to test thoroughly this hypothesis.

In conclusion, our results reveal that CA1 specifically facilitates the retrieval of gist-like memory over at least half a lifetime and receives support from the parahippocampal areas for the most remote memories. In addition, our findings show that CA3 selectively participates to the retrieval of precise memories in a time-limited manner. Using a time-window comparable to that used in human remote memory research allows for our finding to be more readily transferable to humans, hence to further bridge human and animal memory research. Because the nature of the hippocampal contribution to memory retrieval is at the heart of the debates on memory consolidation, our findings have strong implications for current theories of memory consolidation. Indeed, we now show that the nature of memory representations retrieved depends on the functional integrity of specific hippocampal subfields and their time-limited engagement.

## Acknowledgements

We thank Jeannette Maiwald for technical support. This study was supported by a grant from the Deutsche Forschungsgemeinschaft (SA-2146/6-1).

## Author contributions

M.S. and E.A. planned and designed the experiments and wrote the manuscript. E.A. performed behavioural, optogenetic, imaging and histological experiments and analysed the data. S.K and M.T.L performed the electrophysiological recording. All authors approved the final version of the paper.

## Competing interests

The authors declare no competing interests.

## Additional information

**Extended data and Supplementary information are** available

## METHODS

### Animals

One-hundred seventy-two adult male C57BL/6 mice bred at the Institute 8 to 12 weeks old at the beginning of the experiment were included in the analyses. Mice were group housed up to the surgery and immediately afterwards were single housed. Housing conditions were kept under reversed 12 h light/dark cycle (7.00 a.m. light off; 7.00 p.m. light on) to test animals during their active phase. Access to food and water was *ad libitum,* all procedures were approved by an ethics committee of the State of Saxony-Anhalt under license 4252-2 1555 LIN and carried out in accordance with the European Communities Council Directive of September 22^nd^, 2010 (2010/63/EU).

Based on previous studies^1,21^, the sample size was defined as n=4-5 mice per group. Mice were randomly assigned to experimental groups and conditions before starting experiments. Behavioral and imaging data were analyzed off-line by an experimenter blind to experimental conditions. Data collection was performed by an experimenter aware of experimental conditions.

#### Design and general procedures

Different groups of mice and their age-matched controls were tested on a contextual fear conditioning task using retention intervals of 1 day for the recent memory and 6 months or 1 year for the very remote memory. Mice received one footshock when placed in the conditioning context during the encoding phase of the task and were returned to their home cage for the different retentions intervals (one day, 6 months or 1 year; Fig.1A). To test for the precision of the context-footshock association, i.e. to test whether a precise memory or its gist was retrieved with or without CA1 or CA3 cell firing inhibition, mice were re-exposed to the conditioning context and exposed subsequently to a similar but safe context during the retrieval phase of the task (Fig. 1A). Additional experiments were performed to control that exposure to the conditioning context at retrieval prior to exposure to the safe context did not affect the memory performance (% freezing) to the safe context (see Extended Data Fig. 1E) and that baseline exploration of the conditioning and the safe context was comparable for mice that did not receive a shock at encoding (i.e. no-shock mice; Extended Data Fig. 1A,B).To image the neural substrates for the retrieval of the context-footshock association with or without CA1 or CA3 cell firing inhibition, additional groups were tested in the contextual fear conditioning task and immediately sacrificed upon completion of the retrieval phase in the conditioning context to be processed for *Arc* imaging (i.e. assessing memory precision and imaging neural activity induced by retrieving the context-footshock association were not performed within the same animals as the latter requires animals to be to be sacrificed upon completion of the retrieval phase in the conditioning context (Fig 1B) while the former requires animals to be exposed to the similar safe context after being exposed to the conditioning context at retrieval. (Fig.1A). However, behavioral performance of groups used for assessing memory precision and imaging neural activity were comparable (see Supplementary analyses, compare Data Fig 1C-F and Extended Data Fig 3A and B). Cell firing inhibition took place during the retrieval phase of the task.

#### Habituation phase and memory task

Mice were handled for five consecutive days prior conditioning to minimize the effects of stress on the behavioral performance and its neural correlates. Mice underwent surgery for virus/saline injections and optic fiber stubs implantation two weeks before memory for the context-footshock association was tested, consecutively to which they were habituated over 3 consecutive days to be connected to the optical fiber delivering the light stimulation. The fear-conditioning protocol was adapted from the studies of Sauvage et al, 2000 and the Atucha et al., 2015^19,20^. Consecutively to the habituation phase, the conditioning phase of the task started. Mice were randomly allocated to groups prior starting experiments. During the encoding phase, mice belonging to the ‘shock’ groups explored freely the conditioning context for 2.5 min subsequently to which one single footshock (1 mA, 2s) was delivered. Mice remained for another 2.5 min in this context with the goal of consolidating the newly encoded context-footshock association and were returned to their home cage. After a delay of either 1 day, 6 months or 1 year, the memory for this association was tested during the retrieval phase by re-exposing mice to the conditioning context for 5 min (no shock was delivered). Different groups were used for each interval. The precision of the memory for the context-footshock association was tested by subsequently exposing mice to a similar but new context (i.e. to a safe context in which no shock had been delivered; Fig. 1A) also for 5 min. Retrieving the precise memory for the footshock-context association was indicated by a freezing behavior emerging selectively upon re-exposure to the conditioning context (i.e. a lack thereof upon exposure to the safe context). In contrast, retrieving the gist of the memory (i.e. remembering that the footshock was delivered in ‘a’ context but not in which specific context) was indicated by freezing levels comparable in both the conditioning and the safe context. In addition, to identify the neural correlates of precise and gist memory retrieval, additional groups undergoing the same experimental protocol were exposed only to the conditioning context and immediately processed for *Arc* imaging (Fig. 1B; of note, behavioral data for these imaging groups were comparable to those obtained in groups used for assessing memory precision; see Supplementary analyses and Extended Data Fig 3A and B).

The fear-conditioning setup consisted of a 26 X 26 X 38 cm arena with a stainless steel grid floor and four transparent plexiglass walls that constituted the conditioning context. For the safe similar context two opposing walls were covered from the outside with a black and white check-board pattern. A speaker that delivered white noise (55 dB) was attached to the wall. An infrared sensor ring fitted the arena and detected vertical and horizontal activity using the TruScan photobeam activity system (Coulbourn Instruments, Allentown, PA, USA) during the conditioning (encoding) and retrieval phases of the task. Contexts were thoroughly cleaned with 10% ethanol between animals and trials. Behavior of the animals was monitored via video (CCTV, 2.5 MP, IR) to control for gross behavioral abnormalities.

#### Inactivation of CA1 or CA3 during memory retrieval

To transiently and selectively inactivate cell firing in CA1 or CA3 upon memory retrieval in the conditioning and safe context, the inhibitory opsin ArchT was expressed in CA1 or CA3 neurons which led to cell firing inhibition upon blue light LED stimulation. After the five days handling, mice were stereotactically and bilaterally injected in CA1 or CA3 with AAV5.CAMKII.ArchT.GFP.WPRE.SV40 allowing for ArchT expression in pyramidal cells (titer: 1.8e13 [GC/ml], Penn Vector Core Philadelphia, PA, USA; 2 microliters per hemisphere) infused over 10 min with an automated pump as described in Lux et al, 2017^21^.

Histological assessment established that AAV infusions under these conditions were restricted to the targeted areas: CA1 or CA3 (see Extended Data, Fig. 2C and D). Comparable volumes of saline were injected in the saline control groups. Optical Fiber stubs (Plexon, USA) were implanted bilaterally during the same surgery according to Lux et al., 2017^21^. Light stimulation inducing the inhibition of cell firing in the ArchT/Light ON groups started upon placing the mouse in the context (conditioning or safe) and terminated upon completion of the 5 min retrieval phase in either context. The absence of obvious advert structural damage and behavioural effects caused by the light per-se over such a duration had been established in Lux et al, 2017^21^ and was also controlled for in the present study (see saline/ light ON compared to ArchT/light OFF groups in Fig. 1 and results section). Stimulating the inhibitory opsin with a blue LED was chosen over stimulating with a yellow LED because the action spectrum of ArchT extends into the blue range^40,41^ and yellow LEDs commercially available did not provide the sufficient minimal output to induce cell firing inhibition unlike blue light (see Extended Data Fig 2A,B for *In-vivo* electrophysiological evidence of cell firing inhibition with blue LED light stimulation in anesthetized mice, as reported in Lux et al., 2017^21^). Optic Fiber stubs were embedded in a 3D printed mold fitted with a magnet to ease the connection with the optic fiber and minimize stress for animals. Microglass pipets (WVR, USA 30μm tips) were used for AAV injections and kept in place for 1 min after infusion to prevent backflow. Mice were killed at the end of the experiments to assess the location of the optic fiber stubs and the spread of the AAV (see Extended Figure 2C-F and Supplementary Material and Methods). Mice with less than half of CA1 or CA3 infected by the AAV bilaterally or mice for which the AAV could be detected in brain regions bordering the target areas were excluded from the analyses.

#### Surgery

Mice were anesthetized using ketamine 10 mg/kg and xylazine 20 mg/kg, placed into a stereotactic frame (Kopf Instruments,Tujunga, CA, USA) with mouse adaptor (Stoelting, Germany), and the skull was exposed after injecting s.c Carprofen (5mg/kg, Rimadyl A bilateral craniotomy was performed for the implantation of the optical fiber stubs (1.5-mm-long for CA1, 2.00-mm long for CA3; Plexon, USA) and the AAV injections. Surgery was performed 2 weeks before testing to allow for optimal expression of the inhibitory ArchT. The AAV was injected bilaterally at the following coordinates relative to bregma : for CA1: 2 sites: anteroposterior AP: −1.5 mm, mediolateral ML: ±1 mm, dorsoventral DV: −1.5 mm from the surface of the brain and AP: −2.5 mm, ML: ±2 mm, DV: −1.5 mmn. or CA3: 2 sites: AP−1.5 mm, ML ±1,5 mm, DV: −2.0 mm and AP−2.5 mm, ML ±2.5 mm, DV−2.0 mm^31^. Optical fiber stubs were bilaterally implanted at the following coordinates: for CA1: AP:−2.00 mm, ML: ±1.50 mm and for CA3: AP:−2.00 mm, ML: ±1.75 mm. Several layers of dental cement (X200 Auto-mix) were applied and hardened with UV light to anchor the fiber stubs into the skull. A drop of black dye was mixed to the cement to avoid light diffusion. After the surgery, mice were injected with saline (2ml subcutaneous) to prevent dehydration and with metamizol as painkiller (200 mg/kg, for 3 consecutive days) and were given a recovery period of 2 weeks in average which also ensured optimal ArchT expression. Bright field and fluorescent microscopy were used to confirm that the expression of ArchT was restricted to the targeted areas (CA1 or CA3) and that the fiber stubs were appropriately placed (see Extended Data Fig 2C-F and Supplementary Material and Methods).

### Analysis of memory performance

The cumulative time during which an animal did not change its position (resting time) was automatically quantified via an infrared sensor ring detecting vertical and horizontal activity fitting the context arenas using the TruScan photobeam activity system (Coulbourn Instruments, Allentown, PA, USA). Conditioned suppression of activity, e.g. relative resting time or freezing index, was taken as a standard indicator of fear conditioning^1,19,21,39^. Statistical comparisons focused on freezing levels for the first half of the retrieval phase in the conditioning or the safe context (i.e. over the first 150sec). Freezing index was calculated as follows: Freezing index (%) = (resting time of conditioned animals/150)*100. To assess freezing levels specifically tied to the context foot-shock association, freezing levels in conditioned mice were normalized by subtracting freezing levels of age-matched controls that did not receive a footshock, but underwent the exact same behavioral procedure (i.e. the no-shock groups; see Extended Data Fig 1A, B for no-shock freezing levels). Hence, Normalized freezing levels were calculated as follows: Normalized freezing (%)= ((resting time of conditioned animals/150)-(average resting time of no-shock/150))*100. Of note, statistical analyses of non-normalized data yields results comparable to those obtained with normalized data, as was the case in Lux et al, 2016^1^ (see Supplementary Analyses). The absence of effect of the light stimulation *per se* on performance was confirmed by generating a saline/light ON control group. Since this effect was expected to depend on the light stimulation per-se and not on the age of the memory, this group was generated only for the recent memories delay (1 day) in line with the 3Rs principle.

#### Imaging activity in the hippocampus and parahippocampal areas

*Arc* imaging was used to study the neural correlates of the retrieval of precise and gist memories over time and the effects of CA1 and CA3 transient inactivation on these patterns (Fig.2). *Arc* imaging is based on the detection of the RNA of the Immediate-Early-Gene (IEG) *Arc* by in-situ hybridization. This technique yields cellular resolution, hence allows for the percentage of cells recruited during a given event to be evaluated, even in adjacent brain areas such as CA1, CA3 and the parahippocampal areas^1,22^. The detection of the IEG *Arc* was preferred over that of other IEGs such as *Fos* and *zif 268* because it has strong ties to synaptic plasticity, reflects better memory demands and not stress levels^24–26^ and has been heavily used for mapping memory-like activity in the hippocampus and the parahippocampal areas for the past decades. Following the standard protocol described Nakamura and colleagues^25^ and Beer and Vavra and colleagues, 2018^33^, *Arc* pre-mRNA was detected so that only *Arc* intranuclear signal was observable^22,34–36^. In short, mice were a decapitated upon retrieval of the context-footshock association in the conditioning context. Brains were removed, flash frozen in isopentane, and stored at −80°C until sectioning. Brains were sectioned with a cryostat (Leica CM 3050 S; 8-μm–thick coronal sections), mounted on Polylysine slides (Thermo Scientific), and stored at −80°C until *in-situ* hybridization. *Arc* pre-mRNA probes were synthesized using the digoxigenin-labeled UTP kit (Roche Diagnostics). Slides were fixed with 4% buffered paraformaldehyde and rinsed with 0.1 M PBS. Slides were treated with an acetic anhydride/triethanolamine/hydrochloric acid mix, rinsed, and briefly soaked with a prehybridization buffer. The tissue was hybridized with the digoxigenin-labeled Arc probe overnight at +65°C. Following hybridization, slides were rinsed with buffer solutions and treated with an antidigoxigeninhorseradish peroxidase (HRP) conjugate (Roche Molecular Biochemicals) and a cyanin-5 substrate kit (CY5, TSA-Plus system, Perkin Elmer). Nuclei were counterstained with 4’, 6’-diamidino-2-phenylindole (DAPI; Vector Laboratories). Each in-situ hybridization included 1 slide for each group. Slides contained 4 nonconsecutive brain sections (approximately A.P: −3.00 mm and A.P-4.50 mm; Fig 1B)^31^, and images from 3 nonadjacent sections distant approximately 200 microns (i.e., covering approximately 400 microns) at this AP level were acquired. The number of activated neurons was evaluated on approximately 90 neurons per image on 3 nonadjacent sections (i.e., on approximately 270 neurons per area of interest). Images were captured with a Keyence Fluorescence microscope (BZ-X710; Japan). Images were taken with a 40× objective (z-stacks of 0.7-μm–thick pictures; see example Fig.1B. Exposure time and light intensity were kept similar for image acquisition. As first described in the seminal work of Guzowski and colleagues^22^, contrasts were set to optimize the appearance of intranuclear foci^34,37^. To account for stereological considerations, neurons were counted on 8-μm–thick sections that contained 1 layer of cells, and only cells containing whole nuclei were included in the analysis^32^. The quantification of *Arc* expression was performed in the median 60% of the stack in our analysis because this method minimizes the likelihood of taking into consideration partial nuclei and decreases the occurrence of false negative. This method is comparable to an optical dissector technique that reduces sampling errors linked to the inclusion of partial cells into the counts and stereological concerns because variations in cell volumes no longer affect sampling frequencies^32^. Also, as performed in a standard manner in *Arc* imaging studies, counting was performed on cells (>5 μm) thought to be pyramidal neurons or interneurons because small non-neuronal cells such as astrocytes or inhibitory neurons do not express *Arc* following behavioral stimulation^38^. The designation “intranuclear-foci–positive neurons” (*Arc*-positive neurons) was given when the DAPI-labeled nucleus of the presumptive neurons showed 1 or 2 characteristic intense intranuclear areas of fluorescence. DAPI-labeled nuclei that did not contain fluorescent intranuclear foci were counted as “negative” (*Arc*-negative neurons)^22^. Percentage of *Arc-*positive neurons was calculated as follows: *Arc*-positive neurons / (*Arc*-positive neurons + *Arc*-negative neurons) × 100. To focus on the *Arc* signal induced by the retrieval of the context-footshock association, data were normalized (see Fig 2) by calculating the task-induced *Arc* expression by subtracting *Arc* levels observed in “no-shock” groups (see Extended Data Fig. 1A, 1B) from *Arc* levels found in age-matched conditioned mice (i.e. Saline/light ON, ArchT/light OFF, and ArchT/light ON mice) Of note, statistical analyses of non-normalized and normalized data yield comparable results (see Supplementary Analysis and Extended data Figure 5)

#### Statistical analyses

Statistical analyses were performed using SPSS version 27 for windows.

To evaluate the effect of CA1 or CA3 inhibition on memory performance, two- or three-way ANOVAs with freezing levels as dependent variable and the following relevant factors were performed: ‘delay’ (1day, 6 months or 1 year), inhibition (Saline/ArchT-Light ON/ArchT-Light OFF/ ArchT-Light ON), context (Conditioning or Safe) or groups (used for memory precision or for Arc imaging). Within this frame, repeated measures ANOVAs were used when comparing freezing levels between contexts. Two-sided unpaired t-tests were used to compare freezing levels for each area targeted (Saline/ArchT-Light ON/ArchT-Light OFF) and two-sided paired t-tests to compare freezing levels between contexts (conditioning/safe). To assess memory performance one-sample t-tests comparisons to zero were used. To compare the behavioral performance between groups used for memory precision assessment and groups used for imaging neural activity two-way ANOVAs were performed for each target area with ‘freezing levels’ as dependent variable and inhibition (ArchT-Light OFF/ ArchT-Light ON) and delay (1day, 6 months) as factors. For *Arc* imaging, two-way ANOVAs with ‘brain area’ (CA1/CA3/PER/LEC/POR/MEC) and ‘inhibition’ (ArchT-Light OFF/ ArchT-Light ON) as factors and ‘task-induced *Arc* expression’ as the dependent variable were used followed by Tukey post-hoc tests. In addition, one-way ANOVAs with ‘brain area’ as a factor and ‘task-induced *Arc* expression’ as dependent variable followed by Tukey or Dunnet’s posthoc tests were also performed as well as two-sided paired t-tests when relevant. One sample t-tests were used compared to zero to establish the area’s level of engagement. To compare normalized to non-normalized imaging and behavioral data, two-way ANOVAs were performed with ‘*Arc* expression’ or ‘freezing levels’ as dependent variable and ‘inhibition’ (ArchT-Light OFF/ ArchT-Light ON) and ‘delay’ (1day, 6 months) as factors for each targeted area (CA1, CA3).

## EXTENDED DATA AND SUPPLEMENTARY ANALYSIS

*Behavioral (memory) performance was comparable between groups used to evaluate memory precision (Fig 1 C-F) and groups used to assess neural activity (Extended data Fig 3A,B)* as statistical comparisons revealed a lack of group x inhibition interaction effect independent of the hippocampal subfield targeted (recent or very remote memories: for CA1: F_1,12-13_<1.742, P>0.210; for CA3: F_1,12_<0.394; P>0.542).

### Normalizing data did not affect behavioral or imaging outcomes

#### Behavioral data

Statistical comparisons of non-normalized freezing levels (i.e. raw freezing levels; Extended Data Fig.4) yielded results comparable to those obtained with the normalized data. At recent memory tests control groups could discriminate between the conditioning and safe context (Extended Figure 4 A,B; P< 0.027 for paired t-tests, compare white or grey bars between conditioning and safe contexts) indicating that recent memory was also precise in non-normalized data. As reported for normalized data, memory decayed over time and controls retrieved only the gist of the event 6 and 1 year after conditioning as they failed to discriminate between contexts (compare white bars between conditioning and safe contexts: P> 0.385 for all paired t-tests; Extended Figure 4 C–F).

Further statistical comparisons of the non-normalized freezing levels also revealed that, the fear memory was generalized to the safe context upon CA3 inhibition when the memory was recent, i.e. that memory precision was lost under this circumstance as was the case with normalized data (compare green bars between conditioning and safe context: P=0.976 paired t-test; Extended Figure 4A) while very remote gist memory retrieval was unaffected by CA3 inhibition when the memory had decayed 6 months and 1 year after conditioning (compare green and white bars: P> 0.36, unpaired t-tests; Extended Figure 4C, E). Also as shown with normalized data, CA1 inhibition led to impaired retrieval of the gist of the memory when the memory was very remote (compare green and white bars: P < 0.018 unpaired t-tests; Extended Figure 4D,F) and similar results were found at the recent memory test (conditioning context: compare grey or white bar with green bar: P <0.027 unpaired t-tests; Extended Figure 4B).

#### Imaging data

Likewise, analyses of non-normalized and normalized imaging data rendered similar results. Indeed, as it was the case for the normalized data, statistical analyses of the non-normalized data showed that CA3 inhibition was only effective in reducing *Arc* activity in CA3 and no other areas during the retrieval of recent memories (P<0.001 Tukey posthoc tests; F_5,36_=3.09; P=0.020 for area x inhibition interaction effect; Extended data Figure 5A) and did not affect the recruitment of any areas during very remote memory retrieval (F_5,36_=0.247, P=0.938 as area x inhibition interaction effect; Extended Data Figure 5C). This suggests, as was the case with the normalized analysis, that CA3 selectively supports memory precision, and that this contribution depends on the age of the memory trace whereas CA1 contributes to recent gist memory retrieval as CA1 is the only MTL area engaged when the gist of the recent memory is retrieved upon altering of CA3 functional integrity. In addition, as it was the case with the normalized data, the role of CA1 in gist memory retrieval was confirmed in a causal manner as analyses of the non-normalized data showed that CA1 inhibition reduced drastically *Arc* premRNA expression in all areas recruited for the retrieval of gist memories when memory was very remote, (i.e. CA1 and parahippocampal areas: P<0.001 Tukey posthoc tests; CA3: P=0.93; F_5,36_=11.9; P=0.001 for area x inhibition interaction effect; Extended Data Fig. 5D). Likewise, CA1 cell firing inhibition, reduced dramatically activity levels of all brain areas engaged in the retrieval of recent memory (CA1 and CA3: P<0.001; parahippocampus: P>0.856 Tukey posthoc tests; F_5,36_=15.3; P <0.001 area x inhibition interaction effect; Extended Data Figure 5B).

---

**Extended Data Fig 1.**
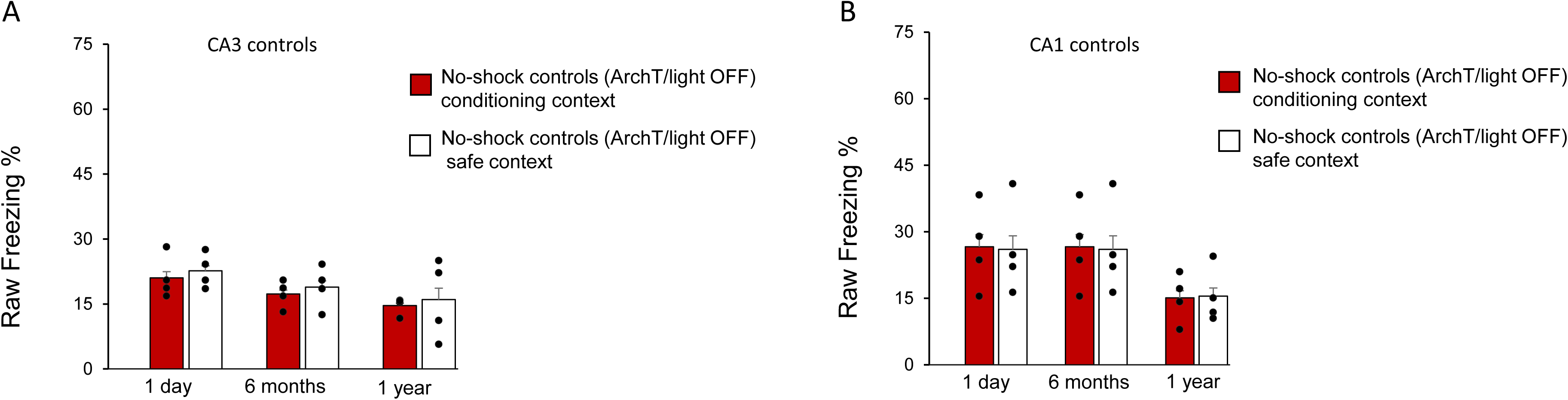
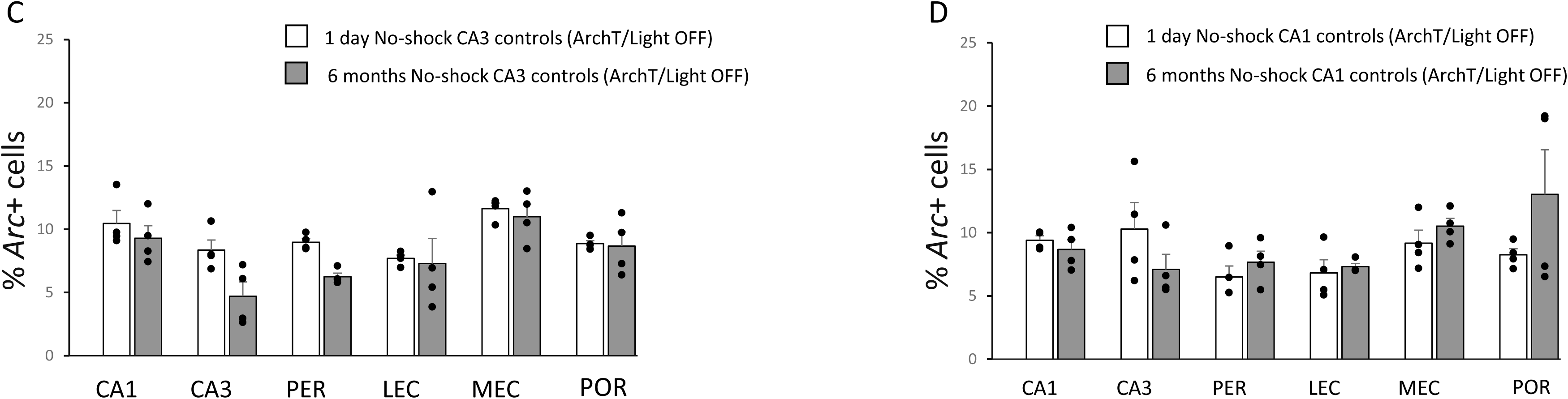
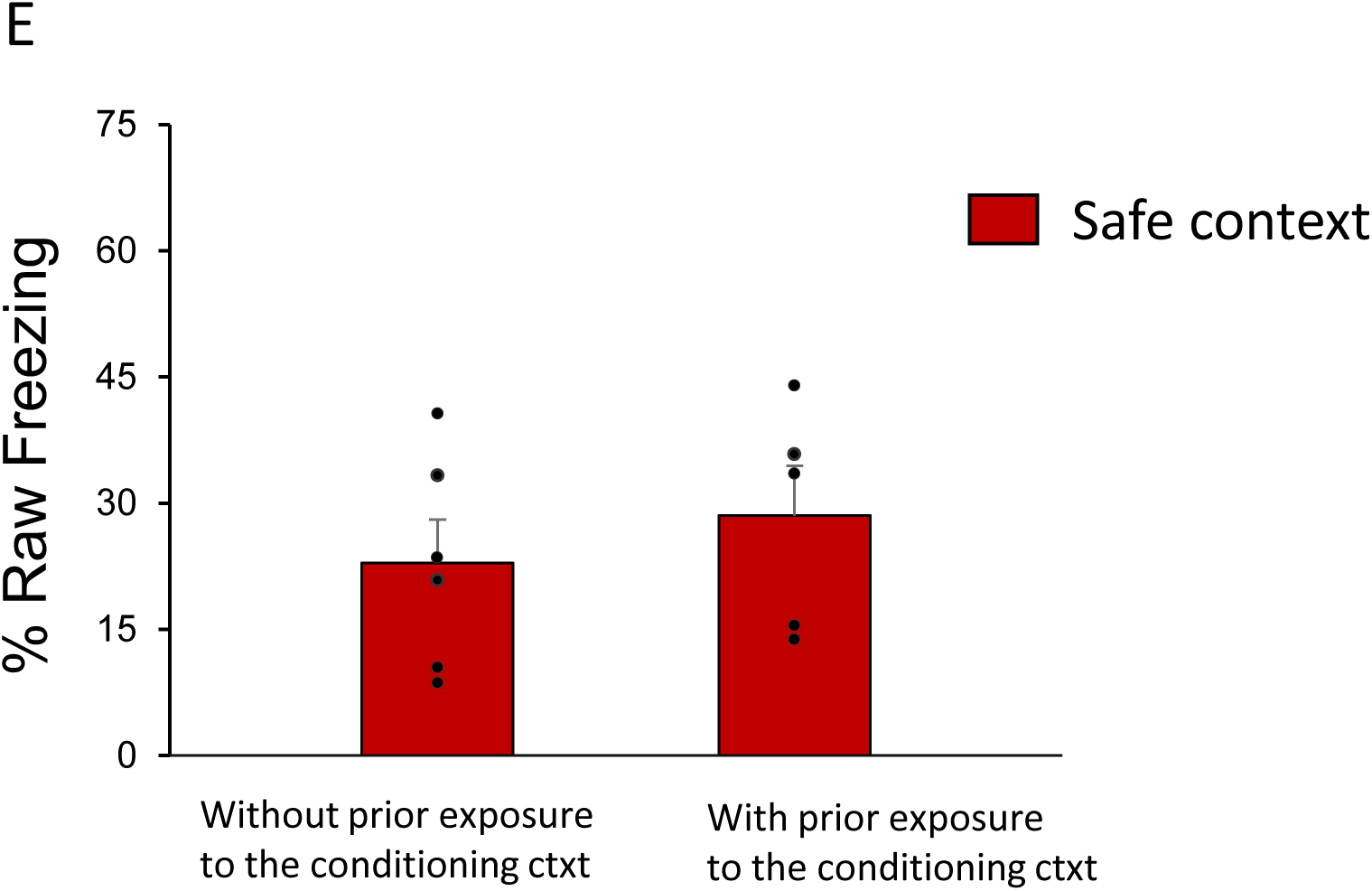
Control measures for task performance and imaging of memory retrieval. *A, B) Raw freezing levels in the conditioning and safe context 1 day, 6 months and 1 year after encoding in control mice that did not receive a footshock at encoding (i.e. no-shock controls).* Mean ± SEM (dot plots overlaid, n=4 per group). Freezing levels did not significantly differ between contexts independently of the age of the memory trace for CA1 or CA3 no-shock controls (F_2,9_<2.53; P>0.135 lack of context x memory age interaction effects with repeated measures ANOVA). Thus, differences in memory performance and brain activity reported upon CA1 or CA3 inhibition in the present study are unlikely to stem from differences in non-cognitive parameters such as locomotor activity or preference/motivation to explore one or the other context. In an attempt at restricting analyses to the freezing levels induced by retrieving the context-footshock association, freezing levels of no-shock controls were used to normalize the freezing levels of ArchT/light ON and ArchT/light OFF conditioned mice by subtracting from their freezing level no-shock controls’ (normalized data displayed in Fig.1). Of note, statistical analyses of normalized and non-normalized (Extended Data Figure 4) freezing levels yielded similar results (see Supplementary analysis). *C, D) Percent of Arc positive cells upon exposure to the conditioning context at test in CA1 D) and CA3 C) no-shock control mice 1 day and 6 months after encoding*. Mean ± SEM. Dot plots overlaid (n=4 per group). As expected in the absence of context-footshock association, neural activity elicited during the retrieval phase of the task was low in both hippocampal and parahippocampal areas (9.48±1.03 % of *Arc* positive cells in average). In addition, neural activity did not significantly differ between recent or very remote memories within area independently of the area targeted CA1 or CA3; Fs_5,36_<1.781, P>0.142, lack of inhibition x delay interaction for each area). This suggests that over time differences in retrieving other aspects of the contextual fear conditioning task than the context-footshock association can unlikely explain the differences in MTL areas engagement over time reported in the present manuscript. To restrict analyses to the brain activity induced by the retrieval of the context-footshock association, neural activity of no-shock controls were used to normalize the level of activity of ArchT/light ON and ArchT/light OFF mice by subtracting from their level no-shock controls’ (normalized data displayed in Fig.2; of note, statistical analyses of normalized data in and non-normalized data displayed Extended Data Figure 5 yield similar results; see Supplementary analysis). CA1: CA1 hippocampal subfield, CA3: CA3 hippocampal subfield, PER: perirhinal, LEC: lateral entorhinal cortex, MEC: medial entorhinal cortex and POR: postrhinal cortex. *E) Raw percent freezing at recent memory test of conditioned mice in the safe context with or without prior exposure to the conditioning context.* Mean values +- SEM. Dot plots overlaid (n=4 per group). Exposing mice to the safe context prior to exposing them to the conditioning context did not affect freezing levels to the safe context (P=0.43, unpaired t-test), indicating that exposure to the conditioning context prior to exposure to the safe context is unlikely to explain the differences in memory performance and neural activity reported in the present study.

**Extended Data Fig. 2:**
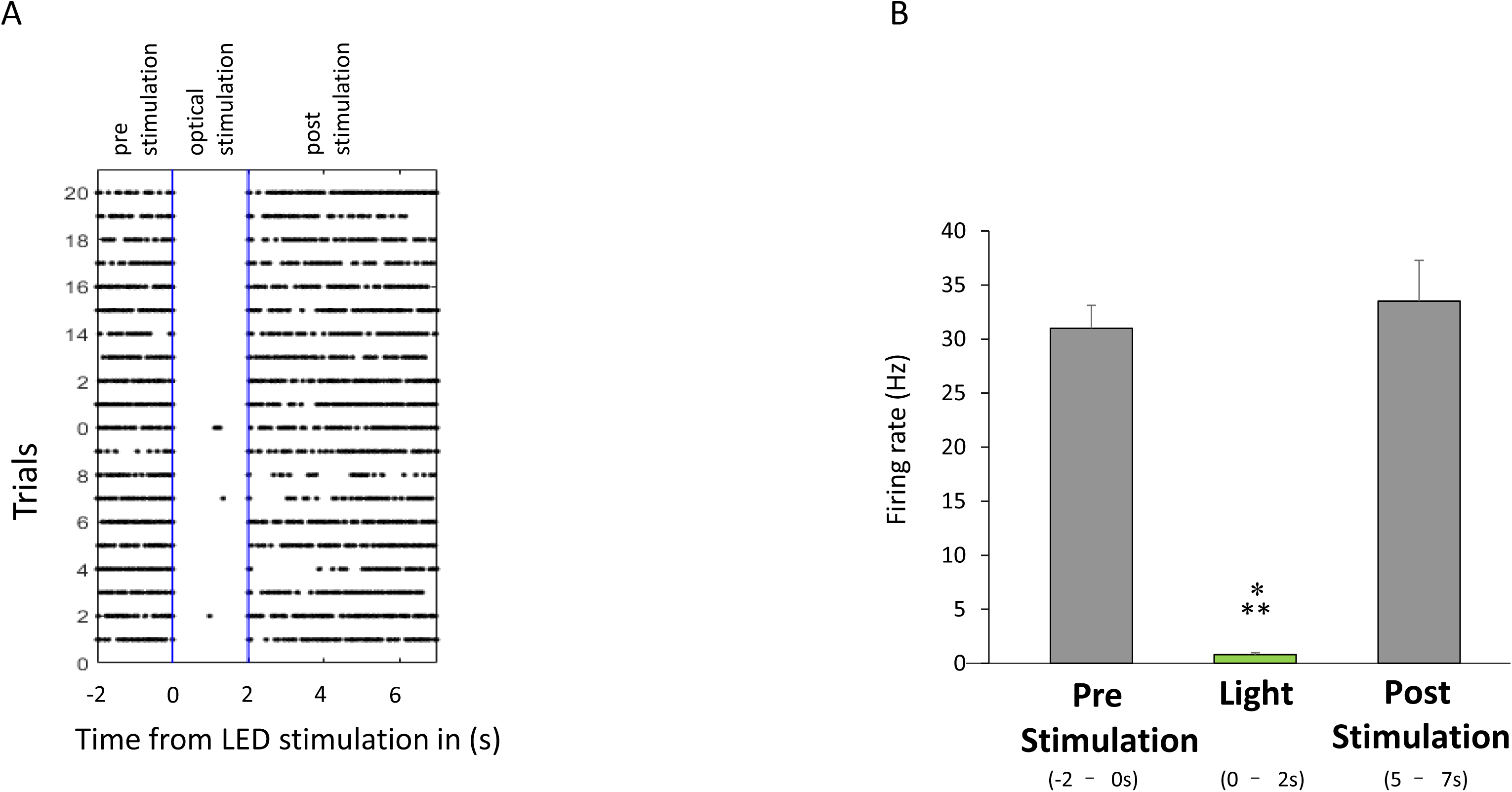
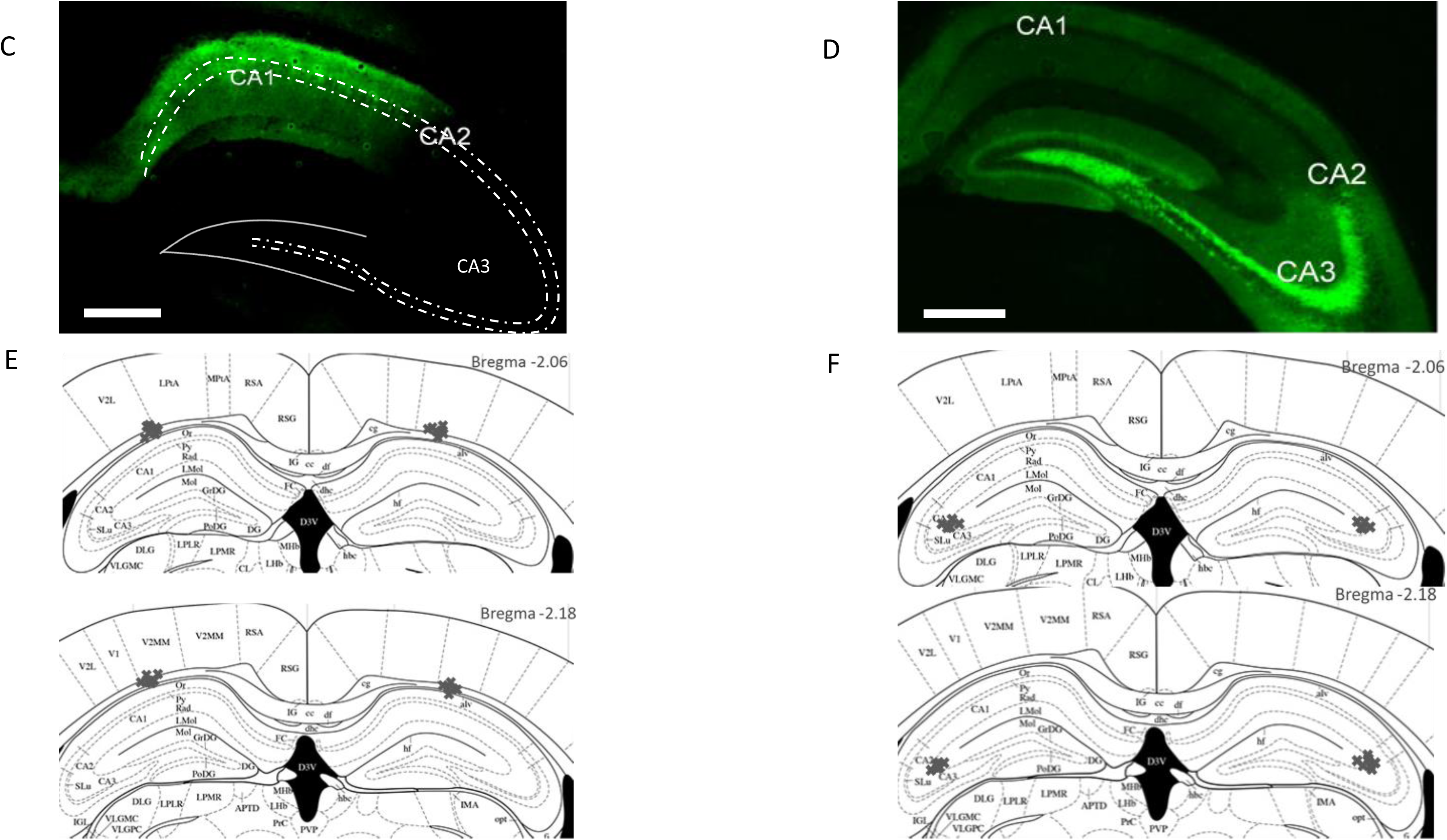
Evidence for inhibition of hippocampal cell firing upon light stimulation *in-vivo*, AAV spread and optical fiber location (see also Supplementary Material and Methods). *A, B) Cell firing is inhibited in the hippocampus upon optical stimulation*. *A)* The raster plot of an optoelectrophysiological recording of an anaesthetized mouse shows that cell firing was abrogated upon light stimulation for each of the 20 trials recorded (9s per trial: 2s pre-stimulation, 2s blue light stimulation, 5 s post-stimulation; 465nm, ∼15-20mW output). *B) Mean firing rates of hippocampal cells* during the 2s pre stimulation, the 2s optical stimulation and the last 2s post-stimulation period. Error bars indicate SEM. Firing rate was drastically decreased upon light stimulation and fully recovered by the last 2s of the post stimulation period (light vs pre- and post-stimulation: P<0.001: pre vs post-stimulation: P=0.862 Tukey post-hoc tests, F_2,57_=55.881, P<0.001 for stimulation effect. *C,D) Histological analysis of the AAV spread on coronal brain sections.* The AAV’s spread was restricted to CA1 (C) or CA3 (D) in all mice included in the analyses. Scale bars, 1mm. *E,F) Diagram of estimated fiber-optic terminations*. Localization of the fiber tips confirmed that all fibers were placed above CA1 for CA1 inhibition (E) and above CA3 for CA3 inhibition (F). No evidence for gross structural abnormalities were observed.

**Extended Data Fig. 3.**
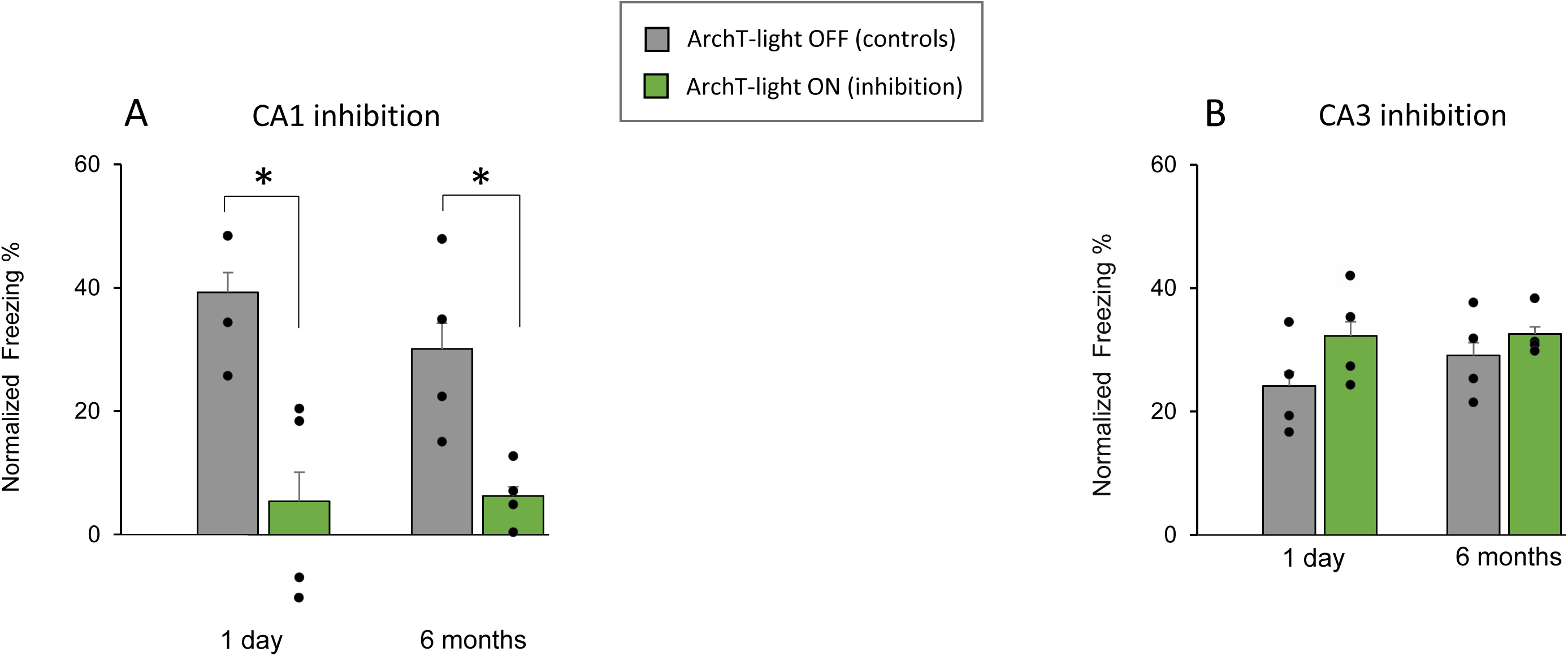
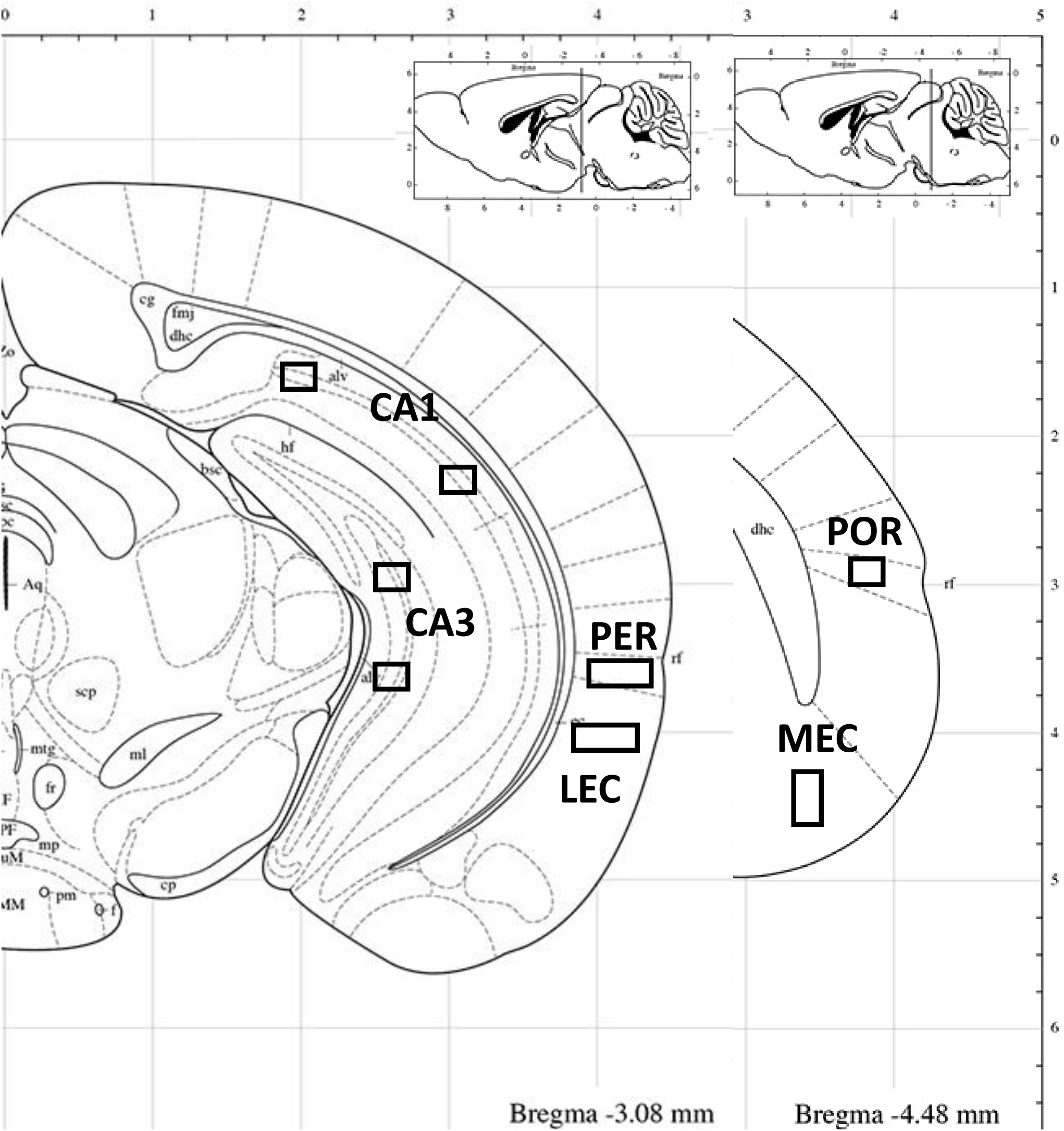
Memory performance of mice undergoing *Arc* imaging and location of the imaging frames. *A,B) Memory performance indicated as mean ± s.e.m., dot plots overlaid.* Normalized percent freezing upon exposure to the conditioning context at test for conditioned animals used to imaging neural activity in the MTL with CA1 A) or CA3 B) as a target. Imaging neural activity induced by the retrieval of the context-footshock association required mice to be sacrificed upon completion of the exposure to the conditioning context during the retrieval phase of the task. This is not compatible with assessing the precision of the memory retrieved as the latter requires exposure to a safe context consecutive to the exposure to the conditioning context. Hence, brain activity elicited by memory retrieval in the conditioning context was imaged in an additional group of mice that underwent the contextual fear conditioning but were sacrificed immediately upon re-exposure to the conditioning context at retrieval as illustrated in Fig 1B. Differences in patterns of activity between recent and very remote memory tests are unlikely to stem from differences in the strength of the memory retrieved one day and 6 months after encoding as memory performance of control groups (ArchT/light OFF) was comparable between recent and very remote memories (compare grey bars between 1day and 6 months: t_6_<1.005, P>0.354, unpaired t-tests; Extended data Fig. 3A&B. As established in groups used to assess memory precision (Fig 1C-H), inhibiting cell firing in CA1 reduced dramatically memory performance independently of the age of the memory (compare grey bars to green bars: P < 0.021 unpaired t-tests) while CA3 inhibition did not affect freezing levels (compare grey bars to green bars:(P < 0.53 unpaired t-tests; see also supplementary analysis for a direct comparison between for performance between memory precision and imaging groups. Of note, analysis of non-normalized data yielded comparable results as shown in the supplementary analysis). *P<0.05. *C) Location of the imaging frames*. To allow for a thorough sampling of the hippocampal subfields CA1 and CA3, CA1 and CA3 frames were taken in the proximal and distal parts of CA1 and CA3 in light of functional differences existing between these levels^25,33,42^. Of note, CA1 and CA3 patterns of activity are not likely to depend on the dorsoventral (DV) or anteroposterior (AP) levels chosen given that patterns of activity in controls in the present study were comparable to those investigated at different AP and DV levels in a previous study of ours that was not focused on identifying the nature of the memory retrieved^1^ in which mice exposed to the same experimental conditions. Parahippocampal frames: PER (perirhinal cortex), LEC (lateral entorhinal cortex), POR (postrhinal cortex) and MEC (medial entorhinal cortex) frames were taken at the same level as previous published work to facilitate comparisons between studies.

**Extended Data Fig. 4.**
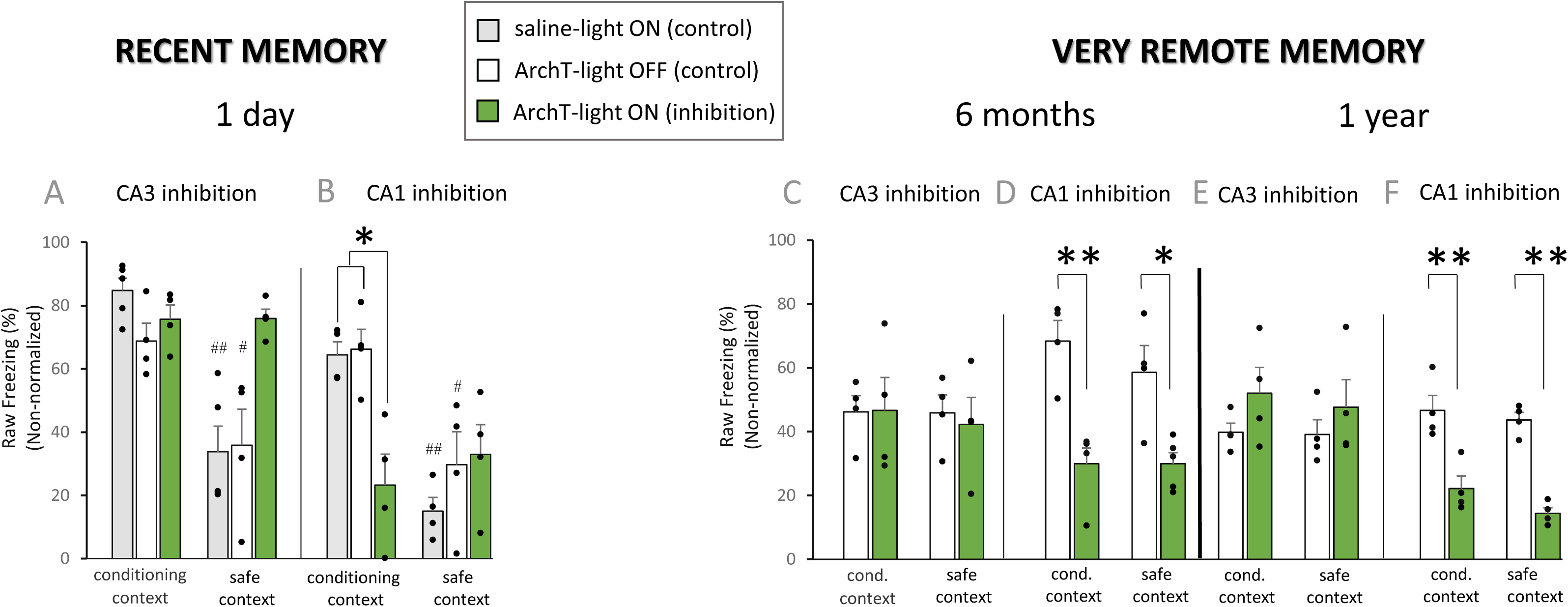
Normalizing behavioural data did not affect outputs. *Non-normalized memory performance at the recent (1 day) and very remote (6 months and 1 year) retrieval tests expressed as mean percent raw freezing ± s.e.m.* (dot plots are overlaid n=4-5 mice per group). To focus on the fear response reflecting the context-footshock association (and not that contributed by other cognitive or non-cognitive parameters), freezing levels of conditioned mice were normalized by levels observed in groups that did not receive a footshock at encoding (i.e. the no-shock groups; see Extended data Figure 1A,B and Material and Methods for normalization) and plotted as in C-H. Normalizing data did not alter the effects observed on memory performance upon CA1 or CA3 inhibition. At recent memory tests control groups could discriminate between the conditioning and safe context (compare white and grey bars between conditioning and safe context; Extended Figure 4A,B; # P< 0.027 for paired t-tests) indicating that recent memory was also precise in non-normalized data. As reported for normalized data, memory decayed over time and controls retrieved the gist of the event 6 months and 1 year after conditioning as they failed to discriminate between contexts at the behavioural level (compare white bars between conditioning and safe context, P> 0.385 on paired t-tests; Extended Figure 4C – F). Statistical comparisons of non-normalized freezing levels revealed that, as was the case with normalized data, the context-footshock association was generalized to the safe context, i.e. memory precision was lost, upon CA3 inhibition when the memory was recent (compare green bars between conditioning and safe context: P=0.976, paired t-test; Extended Figure 4A). In addition, very remote gist memory retrieval was unaffected by CA3 inhibition when the context-footshock memory had decayed 6 months and 1 year after conditioning (P> 0.201, unpaired t-tests, Extended Figure 4C and E). Also as shown with the normalized data, CA1 inhibition impaired the retrieval of the gist of the memory when the memory was very remote (compare white and green bars: P< 0.002, unpaired t-tests for the conditioning context; Extended Figure 4D and F) and similar results were found at the recent memory test (compare grey or white with green bar: P <0.010, unpaired t-tests, Extended Figure 4B). * P<0.05, **P<0.01 for unpaired t-test between green and grey or white bars; # P<0.05, ## P<0.01 for paired t-tests between conditioning and safe contexts (see extended Data Fig. 1 A-E for additional control measures for the task performance).

**Extended Data Fig 5.**
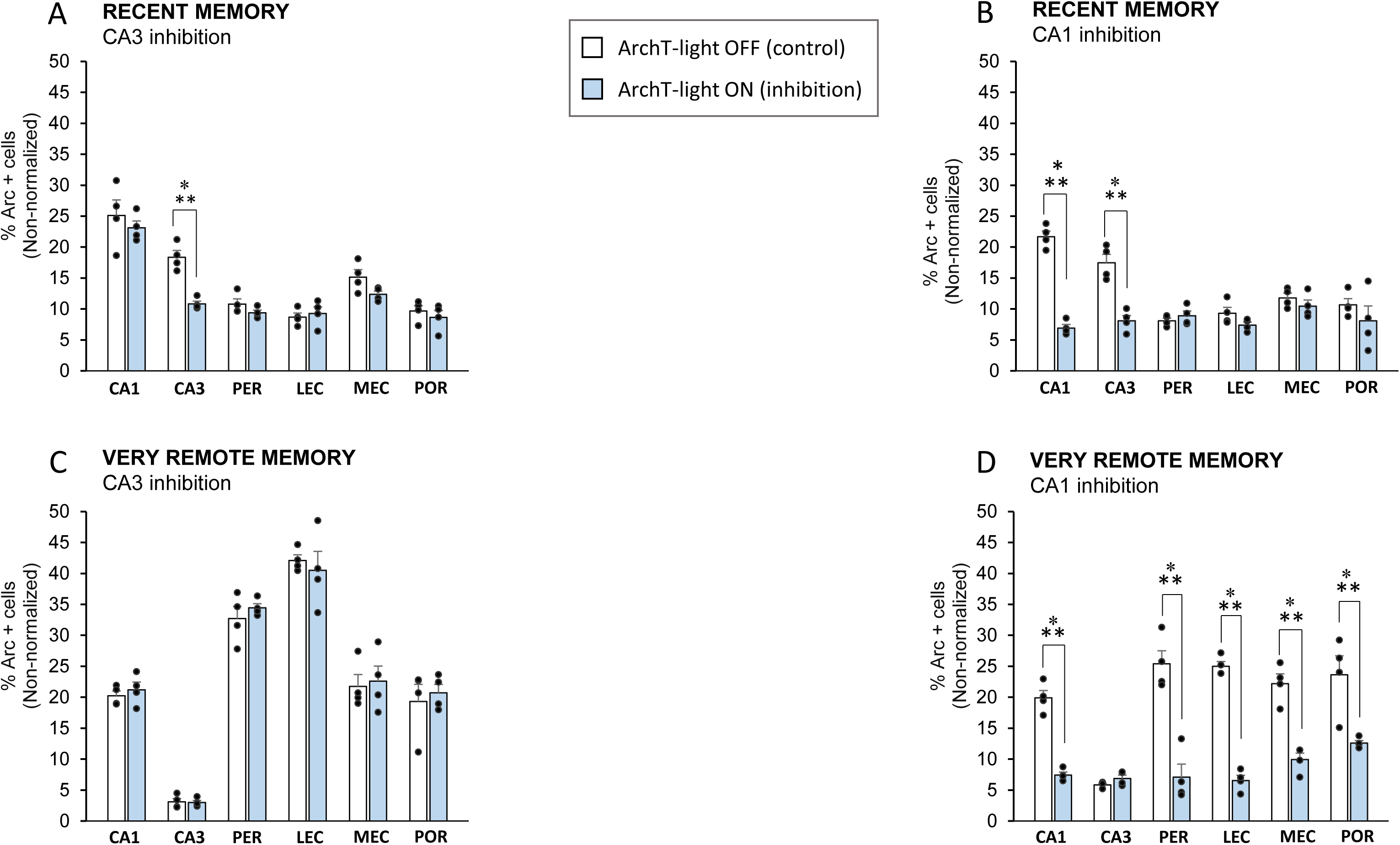
Normalizing Imaging data did not affect outputs. *Neural activity showed as raw percent of Arc positive cells (mean ± s.e.m) of conditioned mice* at the recent (1 day) and very remote (6 months) retrieval tests (dot plots overlaid n=4 mice per group). To focus on the neural activity elicited by the context-footshock association (and not that contributed by other cognitive and non-cognitive parameters), *Arc* levels of conditioned mice were normalized by levels observed in groups that did not receive a footshock at encoding (see Fig. 2 for normalized data, Extended Data Figure 1A,B for no-shock data and Material and Methods for normalization). Normalizing data did not affect changes observed on neural activity upon CA1 or CA3 inhibition. Indeed, as it was the case for the normalized data, statistical analyses of the non-normalized data showed CA3 inhibition was only effective in reducing *Arc* activity in CA3 and no other areas during the retrieval of recent memories (compare white to blue bars: P<0.001 Tukey post-hoc tests, F_5,36_=3.09; P=0.020 inhibition x region interaction effect; Extended data Figure5A) whereas altering CA3 function did not affect the recruitment of any areas during very remote memory retrieval (F_5,36_=0.247, P=0.938 lack of inhibition x region interaction effect; Extended data Figure5C). This suggests, as was the case with the normalized data that CA3 selectively supports memory precision, that this contribution depends on the age of the memory trace and that CA1 contributes to recent gist memory retrieval as CA1 is the only MTL area engaged when the gist of the recent memory is retrieved upon altering the functional integrity of CA3. In addition, as it was the case with the normalized data, the role of CA1 in gist memory retrieval was confirmed in a causal manner as analyses of the non-normalized data showed that CA1 inhibition reduced drastically *Arc* pre-mRNA expression in all areas recruited for the retrieval of gist memories when memory was very remote (compare white and blue bars : CA1 and parahippocampal areas: P<0.001 Tukey post hoc tests; F_5,36_=11.9; P=0.001 for inhibition x region interaction effect; Extended Data Fig. 5D). Likewise, CA1 cell firing inhibition, reduced dramatically activity levels of all brain areas engaged in the retrieval of recent memory (i.e. CA1 and CA3; compare white to blue bars: P<0.001 Tukey post-hoc tests, F_5,36_=15.3; P<0.001 inhibition x region interaction effect Extended Data Figure 5B). ***P<0.001.

## SUPPLEMENTARY MATERIAL AND METHODS

### Evidence for hippocampal cell firing inhibition via ArchT stimulation

To confirm that optogenetic stimulation of the inhibitory opsin ArchT led to cell firing inhibition in the hippocampus, electrophysiological recordings and light stimulation were performed in anesthetized mice injected with AAV5.CAMKII.ArchT.GFP.WPRE.SV40 in CA1 two weeks after injection. Mice were anesthetized (ketamine 10mg/kg and xylazine 20mg/kg i.p) and placed in a stereotactic frame (with a mouse adaptor, Stoelting, Wood Dale, IL, USA). Following a small craniotomy, an optrode was lowered to above the CA1 pyramidal layer. Optrodes were built by gluing a glass-coated tungsten electrode (Alpha-Omega, Israel, 0.5MΩ, shank diameter 125uM, total diameter 250uM) to a 200-μm-thick optical fiber (Plexon Inc., Dallas, Texas, USA) with a maximal tip-to-tip distance of 300 μm. For optical stimulation the optical fiber was connected to an LED driver system (Plexbright, Plexon Inc., Dallas, Texas, USA, 465 nm,∼20 mW output). Broad band neural activity was recorded continuously, with intermittent periods of light application (Neuralynx Digital Lynx SX, Bozeman, MT, USA. Band-pass filter 0.5Hz∼8 kHz, 32 kHz sampling rate). For each neuron, 20 trials were recorded (2s pre-stimulation, 2s light stimulation, 5 sec post stimulation; Extended data Fig. 2 A and B). The optogenetic stimulation and electrophysiology recordings were synchronized through a DAQ system with customized LabView scripts (National Instrument, Austin, Texes, USA). After recording, mice were perfused with 4% paraformaldehyde for histological analysis of the optrode location and AAV spread. For opto-electrical recordings data analysis was performed with customized Matlab codes (Mathworks, 2018). Broad band recordings were high-pass filtered at 300Hz. Multiunit activity was detected by thresholding at 4 standard deviations above baseline.

### AAV spread and location of the optical fiber stubs

The spread of the AAV5.CAMKII.ArchT.GFP.WPRE.SV40 was evaluated on brain sections that had not been processed for *Arc* imaging by detecting its green fluorescent protein (GFP) tag with a Keyence Fluorescence Miscroscope (BZ-X710; Japan) using ×4 and x10 objectives. Animals in which GFP was detected in less than half of CA1 or CA3 bilaterally or in other areas were excluded from the analyses (see Extended Data Figure 2C-D for a representative example of the AAV spread). Sections were also observed under bright field conditions to evaluate fibers placement (Extended Data Figure 2E-F).

## Notes

### Competing Interest Statement

The authors have declared no competing interest.

## REFERENCES

1. Lux, V., Atucha, E., Kitsukawa, T., Sauvage, MM. Imaging a memory trace over half a life-time in the medial temporal lobe reveals a time-limited role of CA3 neurons in retrieval. Elife 11862 (2016).

2. Scoville, WB., Milner, B. Loss of recent memory after bilateral hippocampal lesions. J Neurol Neurosurg Psychiatry 20(1), 11–21 (1957).

3. Anagnostaras, S.G., Maren, S., Fanselow, M.S. Temporally graded retrograde amnesia of contextual fear after hippocampal damage in rats: within-subjects examination. Journal of Neuroscience 19, 1106–1114 (1999).

4. Alvarez, P., Squire L.R. Memory consolidation and the medial temporal lobe: a simple network model. Proceedings of the National Academy of Sciences of the United States of America 91, 7041–7045 (1994).

5. Wiltgen, B.J., Silva, A.J. Memory for context becomes less specific with time. Learning & Memory 14, 313–317 (2007).

6. Winocur, G., Moscovitch, M., Sekeres, M. Memory consolidation or transformation: context manipulation and hippocampal representations of memory. Nature Neuroscience 10, 555–557 (2007).

7. Nadel, L., Moscovitch, M. Memory consolidation, retrograde amnesia and the hippocampal complex. Current Opinion in Neurobiology 7, 217–227 (1997).

8. Frankland, P.W., Bontempi, B. The organization of recent and remote memories. Nature Reviews Neuroscience 6, 119–130 (2005).

9. Yonelinas, A.P., Ranganath, C., Ekstrom, A.D., Wiltgen, B.J. A contextual binding theory of episodic memory: systems consolidation reconsidered. Nat Rev Neurosci 20(6), 364–375 (2019).

10. Broadbent, N. J., Clark, R. E. (2013). Remote context fear conditioning remains hippocampus-dependent irrespective of training protocol, training-surgery interval, lesion size, and lesion method. Neurobiology of learning and memory, 106, 300–308. https://doi.org/10.1016/j.nlm.2013.08.008

11. Bartsch, T., Döring, J., Rohr, A., Jansen, O., Deuschl, G. CA1 neurons in the human hippocampus are critical for autobiographical memory, mental time travel, and autonoetic consciousness. Proceedings of the National Academy of Sciences of the United States of America 108, 17562–17567 (2011).

12. Frankland, P.W., Bontempi, B., Talton, L.E., Kaczmarek, L., Silva, A.J. The involvement of the anterior cingulate cortex in remote contextual fear memory. Science 304(5672), 881–3. (2004).

13. Wheeler, A.L. et al. Identification of a functional connectome for long-term fear memory in mice. PLoS Computational Biology 9, e1002853 (2013).

14. Kitamura, T. et al. Engrams and circuits crucial for systems consolidation of a memory. Science 356(6333), 73–78 (2017).

15. Goshen, I. et al. Dynamics of retrieval strategies for remote memories. Cell 147, 678–689 (2011).

16. Vetere, G. et al. Spine growth in the anterior cingulate cortex is necessary for the consolidation of contextual fear memory. Proc Natl Acad Sci USA 108(20), 8456–60 (2011).

17. Hardt, O., Nader, K., Nadel, L. Decay happens: the role of active forgetting in memory. Trends Cogn Sci 17(3), 111–20 (2013).

18. Rolls, E.T., Treves, A., Foster, D., Perez-Vicente, C. Simulation studies of the CA3 hippocampal subfield modelled as an attractor neural network. Neural Networks 10, 1559–1569 (1997).

19. Sauvage, M.M., Brabet P., Holsboer, F., Bockaert, J., Steckler, T. Mild deficits in mice lacking pituitary adenylate cyclase-activating polypeptide receptor type 1 (pAC1) performing on memory tasks. Molecular Brain Research 84, 79–89 (2000).

20. Atucha, E., Roozendaal, B. The inhibitory avoidance discrimination task to investigate accuracy of memory. Front Behav Neurosci 9(60) (2015).

21. Lux, V., Masseck, O.A., Herlitze, S., Sauvage, M.M. Optogenetic Destabilization of the Memory Trace in CA1: Insights into Reconsolidation and Retrieval Processes. Cereb Cortex 27(1), 841–851 (2017).

22. Guzowski, J.F., McNaughton, B.L., Barnes, C.A., Worley, P.F. Environment-specific expression of the immediate-early gene arc in hippocampal neuronal ensembles. Nature Neuroscience 2, 1120–1124 (1999).

23. Guzowski, J.F., Setlow, B., Wagner, E.K., McGaugh, J.L. Experience-dependent gene expression in the rat hippocampus after spatial learning: a comparison of the immediate-early genes Arc, c-fos, and zif268. J Neurosci 21(14), 5089–98 (2001).

24. Kubik, S., Miyashita, T., Guzowski, J.F. Using immediate-early genes to map hippocampal subregional functions. Learn Mem 14(11), 758–770 (2007).

25. Nakamura, N.H., Flasbeck, V., Maingret, N., Kitsukawa, T., Sauvage, M.M. Proximodistal segregation of nonspatial information in CA3: preferential recruitment of a proximal CA3-distal CA1 network in nonspatial recognition memory. J Neurosci. 33(28), 11506–11514 (2013).

26. Atucha, E., Karew, A., Kitsukawa, T., Sauvage, M.M. Recognition memory: Cellular evidence of a massive contribution of the LEC to familiarity and a lack of involvement of the hippocampal subfields CA1 and CA3. Hippocampus 27(10), 1083–1092 (2017).

27. Sauvage, M.M., Kitsukawa, T., Atucha, E. Single-cell memory trace imaging with immediate-early genes. Journal of Neuroscience Methods 326, 108368 (2019).

28. Wiltgen, B.J. et al. The hippocampus plays a selective role in the retrieval of detailed contextual memories. Current Biology 20, 1336–1344 (2010).

29. Atucha, E. et al. Noradrenergic activation of the basolateral amygdala maintains hippocampus-dependent accuracy of remote memory. Proc Natl Acad Sci USA 114(34), 9176–9181 (2017)

30. Ekstrom AD, Yonelinas AP. Precision, binding, and the hippocampus: Precisely what are we talking about? Neuropsychologia 17(138), 107341 (2020).

## References

31 Paxinos, G., Franklin, K.B. The mouse brain in stereotaxic coordinates. (Gulf Professional Publishing, 2004).

32 West, M.J., Stereological methods for estimating the total number of neurons and synaPes: issues of precision and bias. Trends in Neurosciences 22, 51–61 (1999).

33 Beer, Z. et al. The memory for time and space differentially engages the proximal and distal parts of the hippocampal subfields CA1 and CA3. PLoS Biol 6(8), e2006100 (2018).

34 Vazdarjanova, A., McNaughton, B. L., Barnes, C. A., Worley, P. F., & Guzowski, J. F. (2002). Experience-dependent coincident expression of the effector immediate-early genes arc and Homer 1a in hippocampal and neocortical neuronal networks. The Journal of neuroscience : the official journal of the Society for Neuroscience, 22(23), 10067–10071 (2002)

35 Beer, Z., Chwiesko, C., Kitsukawa, T., Sauvage, M.M. Spatial and stimulus-type tuning in the LEC, MEC, POR, PrC, CA1, and CA3 during spontaneous item recognition memory. Hippocampus 23(12), 1425–38 (2013).

36 Beer, Z., Chwiesko, C., Sauvage, M.M. Processing of spatial and non-spatial information reveals functional homogeneity along the dorso-ventral axis of CA3, but not CA1. Neurobiol Learn Mem 111, 56–64 (2014).

37 Vazdarjanova, A., Guzowski, J.F. Differences in hippocampal neuronal population responses to modifications of an environmental context: evidence for distinct, yet complementary, functions of CA3 and CA1 ensembles. J Neurosci 24(29), 6489–96 (2004).

38 Vazdarjanova, A. et al. Spatial exploration induces ARC, a plasticity-related immediate-early gene, only in calcium/calmodulin-dependent protein kinase II-positive principal excitatory and inhibitory neurons of the rat forebrain. J Comp Neurol 498(3), 317–29 (2006).

39 Rammes, G. et al. Synaptic plasticity in the basolateral amygdala in transgenic mice expressing dominant-negative cAMP response element-binding protein (CREB) in forebrain. Eur J Neurosci 12(7), 2534–46 (2000).

40 Han, X. et al. A high-light sensitivity optical neural silencer: development and application to optogenetic control of non-human primate cortex. Front Syst Neurosci 5, 18 (2011).

41 Chuong, A.S. et al. Noninvasive optical inhibition with a red-shifted microbial rhodopsin. Nat Neurosci 17(8), 1123–9 (2014).

42 Flasbeck, V., Atucha, E., Nakamura, N. H., Yoshida, M., & Sauvage, M. M. Spatial information is preferentially processed by the distal part of CA3: Implication for memory retrieval. Behavioural brain research, 354, 31–38. (2018).

